# *Enterococcus faecalis* delivers Obg GTPase via extracellular vesicles to instigate mTOR activity and promote HCC tumorigenesis

**DOI:** 10.1101/2025.09.30.653999

**Authors:** Ning Ma, Xiaoshan Xie, Jiarui Wang, Zhikai Zheng, Huilin Jin, Xijie Chen, Xiaoling Huang, Haidan Luo, Yue Wei, Qihao Pan, Boyu Zhang, Jiaying Zheng, Peng Zhang, Fenghai Yu, Xue Liu, Zhi-Min Zhang, Zhongguo Zhou, Xiangqi Meng, Mong-Hong Lee

## Abstract

Hepatocellular carcinoma (HCC) is a malignant tumor that has been associated with dysbiosis of the gut microbiota. However, how the gut microbiota plays an oncogenic role in HCC remains largely unknown. Here, we show that *Enterococcus faecalis (E. faecalis)* is highly enriched in liver tumor tissues and is positively correlated with pathogenesis of HCC. *E. faecalis* promotes liver cancer cell proliferation, protein translation, cell migration and tumorigenesis. Mechanistically, we found that EF-derived extracellular vesicles (EF-EVs) deliver EF-Obg GTPase to activate host mTOR, thereby promoting liver cancer progression. Intriguingly, the EF-Obg protein exerts its influence on the mTOR pathway via a Ras-like G domain involved in GTP binding. EF-Obg exhibits a striking homology with the G1 site of the G domain of Rheb, a key positive regulator of mTOR. *Obg* gene is critical for *E. faecalis* to activate mTOR to promote hepatocarcinogenesis based on an engineered *obg* knockdown strain by CRISPR interference experiment. Clinically, abundant EF-Obg protein expression is correlated with enhanced activation of mTOR, leading to poor overall survival in HCC patients. Significantly, treatment of mTOR inhibitor Everolimus confers effectiveness in EF-colonized liver cancer orthotopic model, suggesting that Everolimus therapeutic approach can be effective for liver cancer patients with enriched *E. faecalis*. Taken together, we provide mechanistic and functional evidence to verify a direct causal relationship between tumor-resident *E. faecalis* enrichment and liver carcinogenesis, revealing that EF-Obg functions as a previously unidentified cross-kingdom activator of mTOR to promote liver tumorigenesis.

## Introduction

Liver cancer is one of the top three malignant tumors in the world in terms of mortality rate^1^. Emerging data indicate that gut microbiome could affect liver cancer via gut-liver axis. The destruction of gut barrier, altered bacterial metabolites, and chronic hepatic inflammation are related with HCC progression. Furthermore, tumor tissues of various tumor type, including liver cancer, have a distinct tumor-resident microbiome composition, which are associated with carcinogenesis^2^ and progression^3^. However, the underlying molecular mechanisms of tumor resident microbe-host interaction involved in liver cancer progression remains largely elusive.

*E. faecalis,* a Gram-positive bacterium, is a major cause of hospital-acquired infections^4^. It is also enriched in the gut microbiota of patients with hepatitis C virus-related chronic liver disease^5^, and secretes exotoxin to induce hepatocyte death and liver injury^6^. GelE-positive *E. faecalis* can cause liver tumorigenesis via regulating TLR4-Myd88 signaling^5^. Also, tumor-resident microbiota of liver cancer is increasingly important, but its impacts have not been fully characterized. Tumor-resident *E. faecalis* may employs EVs to influence liver cancer cell biological activities. Mechanistic study such as EVs proteomics, which can reveal functional proteins, may clarify how *E. faecalis*-derived EVs promote liver tumor progression.

The mammalian target of rapamycin (mTOR), a key growth promoter, integrates extracellular signals (nutrition, energy, growth factors) and regulates gene transcription, protein translation, ribosome synthesis, as well as cell growth, apoptosis, autophagy and metabolism^7^. It is estimated that mTOR is frequently activated in 40%-50% HCC cases, reprograming cancer cell metabolism^8, 9^.The classic mTORC1 pathway is activated by growth factors that activate PI3K/AKT and MAPK pathways, which in turn phosphorylate and inactivate TSC1/2, relieving their inhibition on Ras homolog enriched in brain (Rheb). Activated Rheb then activates lysosome-localized mTORC1 and its downstream effectors such as S6K and 4EBP1^10^. Rheb, a Ras family GTPase, can bypass the cell’s response to environmental nutrient levels (such as amino acids) when overexpressed, directly activating the mTORC1 pathway^11^. In yeast, the Rheb homolog protein mediates cellular responses and signal transduction to external nutrient abundance changes^12^. It remains unclear whether tumor-resident bacteria have similar GTPase proteins or other signaling molecules that can regulate the mTORC1 pathway in tumor cells to promote their growth.

In this study, we identified *E. faecalis* as a tumor-resident bacterium that can promote liver cancer. Mechanistically, *E. faecalis* Obg GTPase (EF-Obg) delivered via EVs, which shares homology with Rheb’s G1 site, facilitates the mTOR signaling pathway. CRISPRi-mediated knockdown of *E. faecalis obg* gene impairs its oncogenic role in HCC. High EF-Obg expression correlates with poor overall survival in HCC patients. Importantly, mTOR inhibitor Everolimus significantly suppresses tumor growth in orthotopic liver cancer models exposed to *E. faecalis* administration. Taken together, our studies uncover that *E. faecalis* is an important tumor-resident microbe involved in cancer cell oncogenic signaling pathway activation, verifying a causal relationship between EF-EVs-derived Obg-GTPase and activation of mTOR during liver carcinogenesis. These studies provide insight into diagnosis of HCC exposed to tumor-resident bacterium *E. faecalis* and reveal the therapeutic potential of targeting EF-mTOR axis for HCC with intratumoral *E. faecalis*.

## Methods

### Materials and Methods

#### Ethics Statement

41 fresh-frozen paired samples of liver cancer and para-cancerous tissue, as well as paraffin-embedded liver cancer tissue microarrays (TMA), were collected from the Department of Surgery at Sun Yat-sen University Cancer Center. All samples were collected with the patients’ written informed consent and with approval from the Institutional Review Board of the Sixth Affiliated Hospital of Sun Yat-sen University (ID: G2024006).

All mice were purchased from the Gempharmatech (Guangdong) and were maintained under standard laboratory conditions, with free access to food and water. The animal experiments were conducted in accordance with protocols approved by the Institutional Animal Care and Use Committee of Sun Yat-sen University (No. D2022-0155XS) and the Institutional Animal Care and Use Committee of South China Agricultural University (No. 2023C090).

#### Patient and Public Involvement statement

One hundred patients diagnosed with hepatocellular carcinoma who were treated with radical hepatectomy at Sun Yat-Sen University Cancer Center (SYSUCC) from March 2014 to August 2017 were included. Patients were eligible for the study if they received radical hepatectomy and the samples contained matched tumors and corresponding normal liver tissue. Patients who met the following criteria were excluded: received previous therapies before surgery; received palliative surgery; were diagnosed with other malignant tumors or serious medical diseases; had incomplete medical records or follow-up data.

Clinical data were collected from raw case reports and all of the tumors were staged according to the 8th edition of the American Joint Committee on Cancer (AJCC) TNM staging system and the Barcelona Clinic Liver Cancer (BCLC) staging system. The patients were generally followed up every 3 months in the first 2 years and then every 6 months until recurrence appeared in the following 3 to 5 years. If there was still no recurrence, the patients were followed up once a year. The last follow-up date was July 31, 2024. The overall survival (OS) was defined as the time from the operation date to the death and the recurrence-free survival (RFS) was defined as the time from the operation date to recurrence, or metastasis, or the last follow-up date.

### 16S rRNA gene amplicon sequencing

PCR amplification of the bacterial 16S rRNA genes V3–V4 region was performed using the forward primer 338F (5’-ACTCCTACGGGAGGCAGCA-3’) and the reverse primer 806R (5’-GGACTACHVGGGTWTCTAAT-3’). Sample-specific 7-bp barcodes were incorporated into the primers for multiplex sequencing. The PCR components contained 5 μl of buffer (5 ×), 0.25 μl of Fast pfu DNA Polymerase (5 U/μl), 2 μl (2.5 mM) of dNTPs, 1 μl (10 μM) of each Forward and Reverse primer, 1 μl of DNA Template, and 14.75 μl of ddH_2_O. Thermal cycling consisted of initial denaturation at 98 °C for 5 min, followed by 25 cycles consisting of denaturation at 98 °C for 30 s, annealing at 53 °C for 30 s, and extension at 72 °C for 45 s, with a final extension of 5 min at 72 °C. PCR amplicons were purified with Vazyme VAHTSTM DNA Clean Beads (Vazyme, Nanjing, China) and quantified using the Quant-iT PicoGreen dsDNA Assay Kit (Invitrogen, Carlsbad, CA, USA). After the individual quantification step, amplicons were pooled in equal amounts, and pair-end 2 x 250 bp sequencing was performed using the Illlumina NovaSeq platform with NovaSeq 6000 SP Reagent Kit (500 cycles) at Shanghai Personal Biotechnology Co., Ltd (Shanghai, China).

### Fluorescence in Situ Hybridization (FISH)

The *E. faecalis*-specific probe (5’GGTGTTGTAGCATTTCG) labeled with the fluorophore Cy3 was used to detect the bacterial colonization within human tissues by FISH. The hybridization was performed using a FISH Kit (EXONBIO, D-0016). Briefly, paraffin-embedded sections were de-waxed and hydrated first. The slides were washed in PBS for 10 min twice. Then covered the spots with 0.2 M HCl for 15 min at room temperature. Subsequently, the spots were covered with Proteinase K (50 μg/ml) for 30 min at 37°C and washed with PBS. 20 μl of the hybridization buffer containing the specific probe designed for *E. faecalis* was added. Incubated the slides in a dark chamber for 24 h at 46 °C. After hybridization, the sections were rinsed with wash buffer (20 mM Tris-HCl pH7.4, and 0.9 M NaCl) for 3 times (5 min/time). Air dry the slides for 20 min. 20 μl of DAPI (Life Technologies) were covered and incubated for 15 min at room temperature. The images were captured using Leica Laser Scanning confocal microscope (Leica TCS-SP8, Leica Microsystems Inc, Buffalo Grove, IL, USA).

### Bacteria DNA extraction and qPCR quantification

E.Z.N.A. ® Universal Pathogen Kit (OMEGA, D4035-01) was used to extract bacteria DNA. Briefly, 35 mg tumor tissue samples were added to a Disruptor Tube with 725 μl SLX-Mlus Buffer and vortexed in a TissueLyser (QIAGEN, 85300) with glass beads to lyse and homogenize the samples. The system combined Omega Bio-tek’s MicroElute® LE DNA Columns with RBB Buffer to eliminate PCR inhibiting compounds within the samples and elute highly concentrated DNA. Total genomic DNA was extracted from the tumor tissue and eluted in 100 μl of Elution Buffer, which was preheated to 70 °C. The primers of *E. faecalis* and total bacteria were listed in the Supplemental Table 5. The abundance of the bacterium was calculated as a relative unit normalized to the total bacteria of that sample (where ΔCt=the Ct value of the target (*E. faecalis*) - the Ct value of total bacteria (16S)), using the 2^-ΔCt^ method ^13^.

### Animal models

All mice were purchased from the Gempharmatech (Guangdong) and were maintained under standard laboratory conditions, with free access to food and water. The animal experiments were conducted in accordance with protocols approved by the Institutional Animal Care and Use Committee of Sun Yat-sen University and the Institutional Animal Care and Use Committee of South China Agricultural University.

#### Subcutaneous mouse model

SPF C57BL/6 female mice were pre-treated with an antibiotic cocktail for 2 weeks and then randomly divided into different groups as desired. One week later, indicated materials (*E. faecalis* (1×10^8^ CFU resuspended in 200 μl PBS); EF-CM (200 μl); EF-vec-EVs (50 μg in 200 μl PBS); EF-dCas-Obg-EVs (50 μg in 200 μl PBS); PBS (200 μl)) were administered every other day until the end of the experiment. Mice were subcutaneously injected with 1×10^6^ Hepa1-6 cells one week after the start of gavage. To verify the effects of EF-Obg *in vivo*, C57BL/6 female mice were randomly grouped and then subcutaneously injected with 5 × 10^5^ Hepa1-6 cells infected with plvx-Obg viral selection or plvx-vector viral selection. The tumor length and width were measured every 3 days. The volume was calculated according to the formula (length × width^2^)/2.

#### Orthotopic mouse model

To verify the effects of *E. faecalis* EVs *in vivo*, SPF C57BL/6 female mice were pre-treated with an antibiotic cocktail for 2 weeks and then randomly divided into different groups as desired. One week later, the mice were administrated with indicated treatments (*E. faecalis* EVs (50 ug in 200 ul PBS) or *E. faecalis* (1 x 10^8^ C.F.U resuspend in 200 μl PBS) or the control (200 μl PBS)) every other day until the end of the experiments. 1 × 10^6^ luciferase-expressing Hepa1-6-Luc cells were first injected subcutaneously into SPF C57BL/6 mice. After subcutaneous tumor formation, the tumor-bearing mice were euthanized, and the subcutaneous tumors were harvested. Well-vascularized and viable tumor edges were selected and cut into equal-sized pieces of approximately 1 mm^3^, which were then inoculated into the left lobe of mouse liver one week after the initiation of EVs gavage. Tumor burden was monitored using the IVIS Spectrum In Vivo Imaging System following intraperitoneal injection of D-luciferin (150 mg/kg/mouse). At the end of the experiments, the mice were sacrificed, and their livers were isolated and quantified using an IVIS Spectrum. Quantifications were performed with Living Image v.4.5.2.

### Immunohistochemistry staining

After deparaffinization and rehydration, paraffin-embedded tissue sections (4 μm thick) were treated with retrieval buffer (pH 6.0 sodium citrate buffer) in microwave oven for 10 min. After cooling down, the sections were incubated in 3% H_2_O_2_ for 10 min to quench endogenous peroxidase activity and blocked with goat serum for 1 h. Then the sections were incubated with primary antibody overnight at 4 °C. The slides were then washed with PBS and incubated with secondary antibody at 37 °C for 30 min and stained with DAB substrate. The cell nuclei were stained with hematoxylin. Quantification of the percentages of indicated antibodies per area were performed using ImageJ. Five fields per tumor were chosen for quantification. All primary antibodies used in this study are listed in Supplemental Table 2.

### Cell culture and transfection

HEK293T, Hep3B, SNU449, HUVEC and Hepa1-6 cell lines were obtained from the American Type Culture Collection (ATCC), MHCC-97h cell line was lab storage. These cell lines were maintained at 37 °C and 5% CO_2_ (Thermo, Waltham, MA, USA). SNU449 were maintained in RPMI 1640 medium (RPMI). HEK293T, Hep3B, MHCC-97h, HUVECs and Hepa1-6 cells were maintained in Dulbecco’s modified Eagle’s medium media (DMEM). All the media were supplemented with 10%(v/v) fetal bovine serum (FBS) and 2 × 10^-3^ M L-glutamine. For transient transfection, plasmids were transfected into cell lines followed the standard protocol for Lipofectamine 2000 Transfection Reagent (Thermo Fisher, #11668019).

### Bacteria culture, conditioned medium preparation and EVs isolation

*Enterococcus faecalis* strain ATCC-29212 was obtained from the China General Microbiological Culture Collection Center. *Enterococcus faecalis* strain OG1RF was gifted from Prof Xue Liu’s lab. They were proliferated in brain heart infusion (BHI) medium or BHI agar under aerobic conditions at 37 °C for the rest of the experiments. *E. coli* was grown in Luria-Bertani broth Miller at 37 °C with shaking at 200rpm.

For the conditioned medium preparation, a single *E. faecalis* colony was inoculated into 5 mL of BHI broth for 12 h to reach late exponential phase of growth with OD600nm∼0.8. After centrifugation at 10000 g for 10 min, the supernatant was filtered through Millex-GP Filter Unit (0.22 μm pore size, Millipore). The supernatant after filtration was temporarily stored at 4 °C for further usage^14^.

For the EVs isolation, a single *E. faecalis* colony was inoculated into 5mL of BHI broth and grown overnight. The resulting overnight culture was diluted 1:25 with fresh BHI and grown for 4 hours to reach late exponential phase of growth with OD600nm∼0.8. After centrifugation at 10,000g for 20 min, the supernatant was filtered through Millex-GP Filter Unit (0.22 μm pore size, Millipore). The filtrate was then centrifuged using an ultracentrifugation (Beckman Coulter, USA) at 200,000 g for 2 h at 4°C. The supernatant was discarded, and washed with PBS. The final pellets were suspended with 200 μl PBS and temporarily stored at 4 °C for further usage. EV protein concentration was measured using the BCA method and the particle size and concentration were analyzed by Nanoparticle tracking analysis (NTA).

For the EVs labelling, *E. faecalis* EVs were labelled with 3,3’-dioctadecyloxacarbocyanine perchlorates (DiO, MCE, HY-D0969, 20 μg/ml) for 40 min at 37 °C. The supernatant containing unbound DiO was dia-filtered against PBS buffer using 30 KD Amicon ultra centrifugal filters (Merck Millipore, UFC203024) to remove the unbound DiO dye by centrifugation at 4000 g for 45 min. And protein concentrations were determined. To examine EVs uptake, MHCC-97h grown in 12-well chamber slides were incubated with DiO-labelled EVs or the sham control for different time periods. The slides were then washed with PBS and fixed with 4% PFA for 15 min at room temperature. After washing with PBS, the slides were stained with DAPI for 15 min and imaged with confocal microscopy using a 40X oil immersion objective lens. Fluorescence quantification was performed using ImageJ software.

For the EVs tracing in vivo, EVs were labeled with Cy7 (MCE, HY-D0825) for fluorescent imaging. Mice were oral gavaged with Cy7-labeled EVs and were recorded fluorescence after 1.5 h by IVIS Spectrum In Vivo Imaging System. The mice were sacrificed and harvested livers to analyze fluorescent signals.

### Cell infection

Cell infection by *E. faecalis* was performed as previously described^15^. Briefly, 2 days before infection, cells were seeded in triplicate in 6-well plates. Before infection, the cell culture medium was removed and the cells were washed once with PBS and incubated in serum-free medium for 2 h. *E. faecalis* was harvested and washed twice in PBS, and resuspended in medium without serum to be used at a multiplicity of infection (MOI) of 30. Infection was synchronized by 1 min centrifugation at 1000g. After 3 h of contact, cells were washed 5 times with PBS, and an antibiotic cocktail (150 μg/ml gentamicin and 10 μg/ml vancomycin) was added to kill extracellular bacteria. The efficiency of the antibiotic cocktails was controlled by the absence of viable colonies after plating of the cell supernatants.

### Cell viability assay and colony formation assay

For cell viability assay, cells were seeded into 96-well plates at a density of 1000 cells per well. 10%(v/v) Cell Counting Kit-8 (K1080, Apexbio Technology LLC, Houston, TX, USA) solution was added to the well. After 3 h incubation, optical density at 450nm was measured using EpochTM Multi-Volume Spectrophotometer and Take3TM (BioTek, Winooski, VT, USA).

For colony formation assay, cells were seeded into 6-well plates at a density of 800 cells per well. After 7 to 10 days, cells were washed three times with PBS, the colonies were fixed with 4% polyoxymethylene and stained with 0.05% crystal violet at room temperature for 30 min. Number of colonies was calculated by Image J software.

### Tube formation assay and migration assay

For the tube formation assay, matrigel matrix (Corning, USA) was first thawed at 4°C and spread into 96-well plates with 50 μl in each well and rested at 37 °C for 30 min to form a gel. 1×10^4^ pretreated (co-cultured with *E. faecali*s at MOI of 30 for 3 h, or 10% EF-CM for 24 h, or 50 μg/ml EVs for 24 h) HUVEC were suspended in 100 μl of the culture medium. After 16 h incubation, tube formation was photographed with a digital camera system, and the total length and number of branches of the tubes were analyzed with Image J software.

For the migration assay, pretreated 5 × 10^4^ HUVEC cell suspended in 200μm medium without FBS were seeded in the upper chamber (diameter = 8 μm; Corning Costar, New York, USA), and 600 μl medium containing 10% FBS were added to the lower chamber. After 24 h of incubation, cells were stained with 0.1% crystal violet for 30 min and 5 random fields of view were selected to count the number of migrating cells. Photographs were taken for counting with a digital camera system.

### RNA Seq Analysis

RNA-Seq analysis was performed in Hep3B cells with or without infection with *E. faecalis* and 10% EFCM treatment. Total RNA extracted from the indicated groups of Hep3B cells was subjected to RNA-Seq performed by SHANGHAI BIOTECHNOLOGY CORPORATION (Shanghai, China). The sequencing reads were analyzed to obtain expression profiles. Gene set enrichment analysis (GSEA) was performed used the GSEA software provided by the Broad Institute (http://www.broadinstitute.org/gsea/index.jsp), in accordance with the instructions provided by the Broad Institute.

### Immunoblotting

The immunoblotting was conducted as previously described^16, 17^. Total cell or tissue lysates were solubilized in lysis buffer (50 mM Tris–HCl, 1 mM EDTA, 150 mM NaCl, 0.1% NP-40, 0.1% Triton-100) containing protease inhibitors cocktail and phosphatase inhibitors (Bimake). Lysates were immunoprecipitated with indicated antibodies. Proteins were resolved by SDS-PAGE gels and then proteins were transferred to polyvinylidene difluoride membranes (Millipore). The membranes were blocked with 5% nonfat milk for

1 h at room temperature followed by incubation with indicated primary antibodies. Subsequently, membranes were washed in Tris-buffered saline with Tween-20 (Sangon Biotech, #A600669-0250) there times and incubated with indicated peroxidase-conjugated secondary antibodies (Thermo Scientific, #31430) for 1 hour at room temperature. Following several washes, chemiluminescent images of immunodetected bands on the membranes were recorded on X-ray films using the enhanced chemiluminescence (ECL) system (Bio Rad, #170-5061). The primary antibodies are listed in Supplemental Table 2.

### Anti-puromycin immunoblot analysis

Cells were plated in pre-warmed culture medium in sterile tissue culture plates and treated with the indicated treatment. After completing the treatments, the cells were washed twice with pre-warmed sterile PBS, and were cultured with 1 μM puromycin under normal culture conditions for 30 min. At the end of the incubation period, medium containing puromycin was aspirated, and the cells were washed three times with ice-cold PBS. The cells were then harvested to extract proteins for immunoblot assays. The PVDF membrane was incubated with anti-puromycin antibody.

### RNA isolation and quantitative real-time PCR

Total RNA was extracted from tissues and cells using TRIzol reagent (Invitrogen) and 1μg RNA was reverse transcribed to complementary DNA (cDNA) using ReverTra Ace® qPCR RT Master Mix with gDNA Remover (TOYOBO, Osaka, Japan) according to the manufacturer’s instructions. Quantitative real-time PCR analyses were performed using 2 × SYBR Green qPCR Master Mix (Bimake) and specific primers on a LightCycler 480 (Roche, Basel, Switzerland). All sequences for qRT-PCR used in this study are listed in Supplemental Table 5.

### Plasmids Construction and overexpression cell lines construction

*Obg* cDNA was amplified by PCR from *E. faecalis* cells and cloned into PLVX vector and PCDNA3.1 with HA and MYC tag. The primers were listed in the Supplemental Table 3. For lentivirus preparation, HEK293T cells were seeded in a 10cm dish at a density around 1 × 10^6^, and were co-transfected with 10 μg PLVX-Obg plasmid, 5 μg psPAX2 and 5μg pMD2.G by using polyethyleneimine (Polysciences, 24765). The supernatant which contained lentivirus was collected at 48 and 72 h after transfection, and were filtered through Millex-GP Filter Unit (0.22 μm pore size, Millipore). Hepa1-6 cells were infected with filtered viral supernatant containing 10 μg mL^−1^ polybrene (Millipore, TR-1003-G), followed by puromycin selection and finally verified by western blot. PCDNA3.1-Flag-mTOR was kept in the lab.

### Obg overexpression in *E. coli*

Obg cDNA was amplified by PCR from *E. faecalis* cells. The products were digested with HindIII and BamHI restriction enzymes, ligated into a PET21a vector using T4 DNA ligase, and transformed separately in DH5a cells of *E. coli* for cloning. Colonies on ampicillin agar plates were screened for successful construct formation by colony PCR and double restriction digestion. The recombinant constructs were transformed into BL21 (DE3) competent cells of *E. coli*, and protein expression was induced with 1mM isopropyl β-D-1-thiogalactopyranoside (IPTG) at 18 °C for 16 h. The cells were isolated by centrifugation (6000 g, 4°C), dissolved in lysis buffer (50 mM Tris–HCl pH 7.5, 1 mM EDTA, 150 mM NaCl, 0.1% NP-40, 0.1% Triton-100, 0.01% SDS) containing protease inhibitors cocktail and phosphatase inhibitors (Bimake), and disrupted by sonication. The supernatant-containing proteins were collected after centrifugation (12000 g, 4 °C, 30 min). The primers were listed in the Supplemental Table 3.

### Coimmunoprecipitation (Co-IP)

The coimmunoprecipitation was conducted as previously described^18, 19^. After indicated treatment, cells were lysed with cell lysis buffer (50 mM Tris-HCl pH 7.5, 1 mM EDTA, 150 mM NaCl, 0.1% NP-40, 0.1% Triton-100) containing protease inhibitors cocktail and phosphatase inhibitors (Bimake). For each lysate, supernatants were collected after centrifugation and incubated with appropriate antibodies overnight at 4°C, followed by Anti-Myc Magnetic Beads incubation for 4 h. After incubation, beads were washed with cell lysis buffer for three times. After that, add propriate volume of 2 × loading buffer to the beads and boiled for 10 min in 95 °C to get the eluted proteins. Next, immunoblot assays were performed with specific antibodies The primary antibodies are listed in Supplemental Table 2.

### Preparation of Obg antibody

The anti-Obg antibody was prepared by SinoBiological Confidential in Peking. 5 Balb/c mice were immunized each with two VLP-conjugated peptides. After four immunizations with the protein and three with the peptides, ELISA binding assays confirmed that all 10 mice met the fusion criteria for immunogenicity. After 1 hybridoma fusion, a total of 2 strains of hybridoma cell strains were obtained that bound to both antigenic proteins and clinical samples provided by the client. Subsequently, 4 strains of mouse monoclonal antibody were purified to meet the required quantity and achieved purity exceeding 90%. These antibodies were validated to bind to the antigen proteins in ELISA assays.

### In situ proximity ligation assay (PLA)

The standard commercial protocol (Sigma-Aldrich, DUO92101) was followed for conducting PLA ^20^. The fixed HKT293T cells were subjected to permeabilizing in a 0.5% Triton X-100 solution for 10 min. Subsequently, PLA blocking solution was employed for blocking for 1 h prior to the incubation of the primary antibodies. After overnight incubation in a humidified chamber at 4 °C, the samples were treated with PLA secondary probe and incubated at 37 °C for 1 h. The ligation process was completed by applying Ligation mix to each sample, incubating them at 37 °C for 30 min. Polymerization mix was used for amplification and the samples were further incubated at 37 °C for 100 min. After incubation, 1 × Buffer B was used to wash the samples once for 10 min, followed by another wash with 0.01 × buffer B for 1 min at room temperature. Finally, the samples were prepared for imaging by mounting them with Duolink in situ mounting medium containing DAPI for 15 min. The proximity ligation signal was captured using a confocal microscope.

### *Obg* Knockdown by CRISPR interference system

The CRISPR interference was performed as previously described^21^. Briefly, two plasmids were used: pMSP3545-dCas9str (Addgene, #153516) encoding dCas9, and pGCP123-sgRNA encoding sgRNA. The sgRNA targeting the gene *obg* was designed on the CHOPCHOP website (https://chopchop.cbu.uib.no/) The sequence was synthesized by BGI (The Beijing Genomics Institute) and cloned into plasmid pGCP123-EbpA-g1(Addgene, #153517) with restriction sites PstI and BamHI, and ligated with T4 DNA ligase. This plasmid was subsequently transformed into DH5a. The sgRNA target *obg* sequence is listed in Supplemental Table 4.

As for the preparation of *E. faecalis* electrocompetent cells, we followed the procedure provided by Dr. Kimberly A Kline. Briefly, *E. faecalis* dilute culture 1:10 into prewarmed fresh BHI (100 ml) and incubate until culture OD at 600nm reaches 0.5-0.75. Then chill cells on ice and pellet cells at 6000 rpm for 10 min at 4 °C. Resuspend pellet in 2 ml ice cold sterile H_2_O and split into two 1.5 ml tubes. Pellet at 13000 rpm for 1 min. Resuspend each pellet in 500 μl lysozyme solution (10 mM Tris pH 8.0, 20% sucrose, 10 mM EDTA, 50 mM NaCl) containing 30 μg/ml lysozyme. Incubated at 37°C for 20 min. Wash each pellet 4 times with 0.5 ml ice-cold electroporation buffer (0.5 M Sucrose, 10% glycerol). Resuspend each tube in a total of 300 μl EB/tube and split into 50-65 μl aliquots. Use fresh or store at -80°C.

Electroporation was performed in 0.2-cm-gap electroporation cuvettes (Bio-Rad) at 25μF, 400Ω, and 2.5kV. After electroporation, 1ml SBHI medium, containing 0.5M sucrose in BHI medium, was immediately added to transformed cells, and the cells were left to recover for at least 2 h at 37°C without shaking. The recovered cells were then plated on selective BHI agar plates. Erythromycin (2 μg/ml) was used to maintain the pMSP3545 plasmid in *E. faecalis*, and kanamycin (500 μg/ml) was used to maintain pGCP123 and its derivatives. We induced the target gene expression with 25 ng/ml nisin (MCE, HY-P1607). The inhibition efficiency of Obg was evaluated by qPCR.

### Statistical analysis

Mean ± SD (at least three biological replicates or three independent experiments) was used to plot the data. Student’s *t*-test, one-way Analysis of Variance (ANOVA) test, and two-way ANOVA test were used to analyze quantitative data between groups. Kaplan‒ Meier curves were generated to estimate OS, RFS, and differences between curves were evaluated using a log rank test. A value of p < 0.05 was considered statistically significant. *, *p* < 0.05; **, *p* < 0.01; ***, *p* < 0.001; ****, *p* < 0.0001; ns, not significant;

## Results

### *E. faecalis* is enriched in liver cancer tissue and can promote liver cancer cell proliferation and migration

To investigate the contribution of microbes colonizing in liver tissue during the progression of hepatocellular carcinoma (HCC), the microbiota composition of tumor tissues and para-cancer tissues (as normal tissues) was profiled by 16S rRNA sequencing. Although the principal coordinate analysis (PCoA) showed no significant compositional differences (Figure S1A and S1B), alpha diversity was reduced in liver tumor tissues (Figure 1A). Additionally, *Firmicutes_D* abundance positively correlated with HCC biomarker Des-gamma-carboxyprothrombin (PIVKA_II); *Enterococcaceae* (under *Firmicutes_D*) positively correlated with PIVKA_II and Aspartate Aminotransferase (AST) levels (Figure 1B and 1C), suggesting *Enterococcaceae* may promote hepatocarcinogenesis.

**Figure 1.**
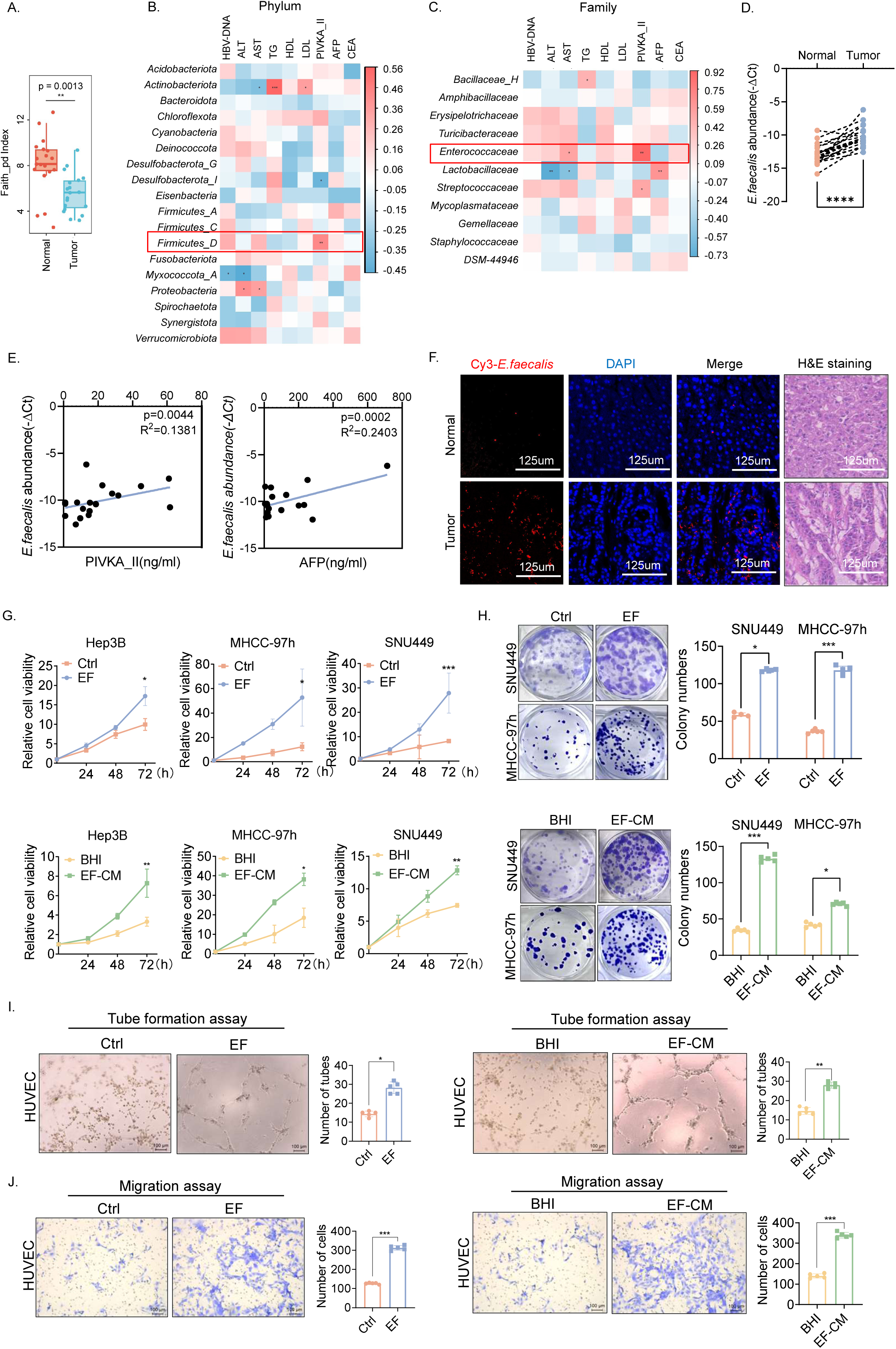
*E. faecalis* is enriched in liver cancer tissue and promotes proliferation of liver cancer cell. **(A)** Faith_pd index showed the alpha diversity of microbiota extracted from liver tumor tissues and normal tissues (n = 19). **(B&C)** Spearman’s correlations between the relative abundance of bacterium and clinical parameters of HCC patients. The red box indicates the Phylum and Family of *E. faecalis*. **(D)** Relative *E. faecalis* abundance in tissues detected by qPCR. **(E)** Spearman’s correlation between *E. faecalis* abundance and serum PIVKA_II or AFP level in HCC patients. **(F)** Detection of *E. faecalis* in tissues by fluorescence in situ hybridization (FISH). H&E staining was shown. Scale bar = 125 μm. **(G&H)** CCK8 assay and Colony formation assay of indicated HCC cells. **(I&J)** Tube formation and migration assay of HUVEC cells were shown. Scale bar = 100 μm. Wilcoxon test was used to calculate the *p* value between 2 groups (A). Unpaired two-tailed Student’s *t* test were used in (B), (C), (G), (H), (I) and (J). Paired *t*-test was used in (D).

To further define the specific species of *Enterococcaceae* associated with HCC, we quantified the abundance of several well-studied species, *E. casseliflavus, E. faecium, E. faecalis* and *E. mundtii* within tissue samples (n=40) using real-time PCR. The results showed that *E. faecium* and *E. mundtii* were almost undetectable in all samples. *E. faecalis* was highly enriched in tumor tissues compared with normal tissues (Figure 1D), while *E. casseliflavus* showed no significant difference (Figure S1C). Correlation analysis revealed *E. faecalis* (but not *E. casseliflavus*) abundance significantly correlated with PIVKA_II and AFP levels (Figure 1E and S1D). Tumor-resident *E. faecalis* colonization was confirmed via FISH (Figure 1F), suggesting *E. faecalis* is a tumor-enriched microbe correlated with HCC clinical parameters.

To determine *E. faecalis’* oncogenic potential in HCC, we cocultured HCC cell lines with *E. faecalis* (MOI 30). *E. faecalis* promoted HCC cell proliferation and colony formation (Figure 1G and 1H). It also enhanced HUVEC tube formation and migration (Figure 1I and 1J). Since bacterial metabolites and secreted proteins may play a significant role in these phenomena^14^, we prepared the conditioned medium (hereinafter referred to as EF-CM) for examining impacts on the growth/migration. Indeed, EF-CM also promotes proliferation of HCC cells and tube formation and migration of HUVEC cells (Figure 1G to 1J). These results indicate that *E. faecalis* and its conditioned medium can promote HCC cell growth and HUVEC cell migration.

### *E. faecalis* potentiates mTOR signaling pathway and promotes HCC tumorigenesis

To elucidate the precise mechanism, we conducted RNA sequencing analysis on Hep3B cells. Gene Set Enrichment Analysis (GSEA) revealed that mTOR pathway was enriched in both groups (Figure 2A and 2B). Heatmap results showed upregulation of most mTOR downstream genes after indicated treatment (Figure 2C). Both *E. faecalis* and EF-CM enhanced phosphorylation of mTOR and its downstream effectors P70S6K and 4EBP1 (Figure 2D). Additionally, anti-puromycin immunoblot analysis confirmed increased protein translational activity following the indicated treatments (Figure 2E). Collectively, these findings suggest that *E. faecalis* and EF-CM potentiates mTOR signaling pathway, including enhancing protein synthesis.

**Figure 2.**
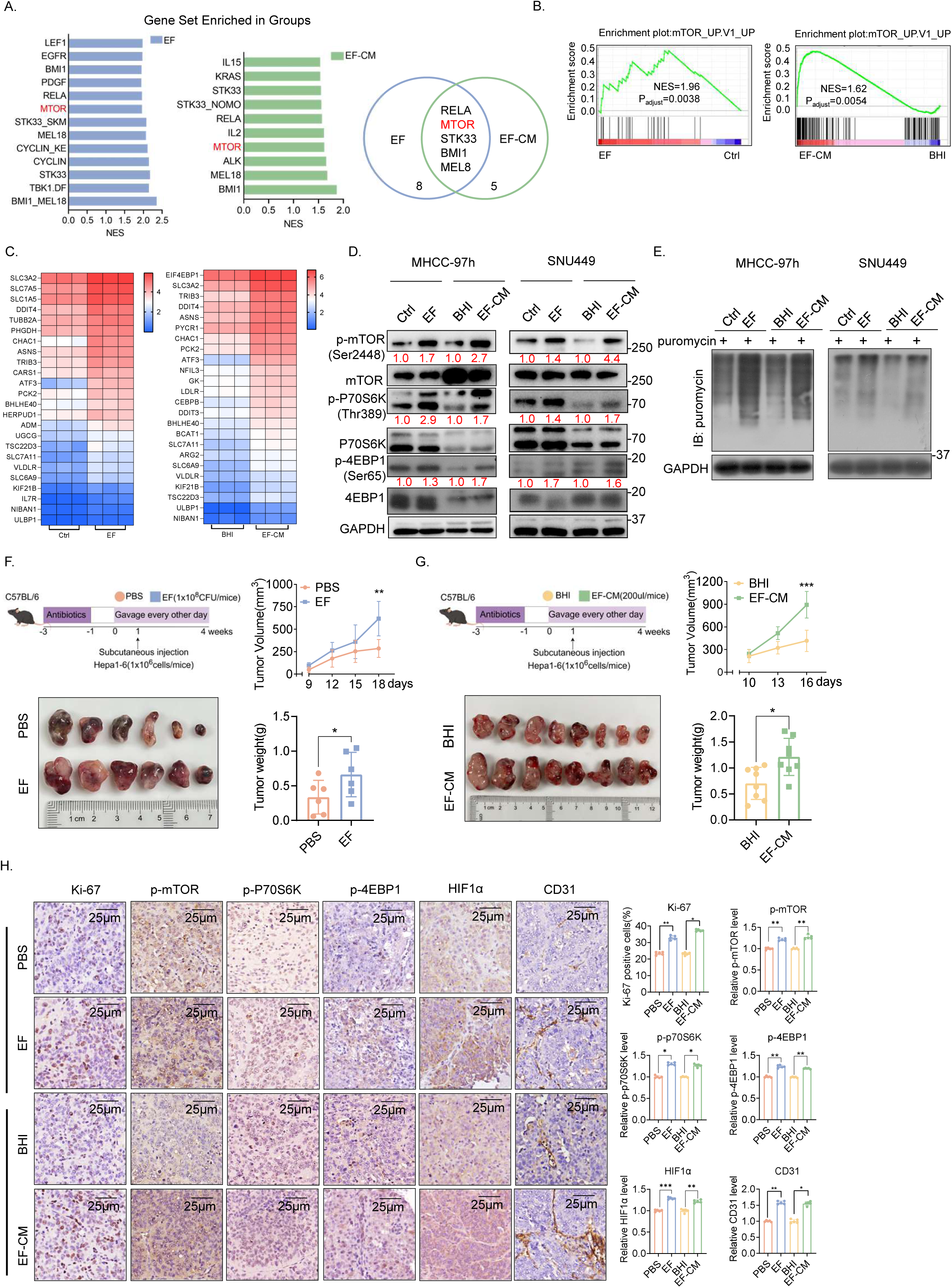
*E. faecalis* enhances mTOR signaling activity and promotes liver cancer tumorigenesis. **(A)** Significantly enriched cancer-related pathways in Hep3B cells after indicated treatment. Overlapping pathways enriched in both EF and EF-CM treatment (NES > 1.5) were shown. **(B)** GSEA plots of mTOR signaling pathway correlated with indicated treatment. **(C)** Heatmap showed mRNA levels of mTOR signaling pathway-related genes after indicated treatments, as revealed by RNA-sequencing analysis. **(D)** Protein levels of mTOR, P70S6K, 4EBP1 and their phosphorylation levels in HCC cells. **(E)** Anti-puromycin immunoblot analysis showed translation activities in HCC cells. **(F&G)** Schematic diagram of animal experiments. Tumor volumes, tumor weights and representative tumor pictures were shown (n = 6 in F, n = 8 in G). **(H)** Representative images of immunohistochemical staining from tumor samples. Permutation test was used to calculate the *p*-value in (A) and (B). Unpaired two-tailed Student’s *t* test was used in (F), (G) and (H). NES, Normalized enrichment score.

Furthermore, we conducted a subcutaneous allograft experiment in C57BL/6 mice using mouse liver cancer cells Hepa1-6. *E. faecalis* or EF-CM (Figure 2F and 2G) was administered intragastrically every other day, PBS or BHI was administered as control respectively. The results showed that administration of *E. faecalis* or EF-CM promoted tumor growth (Figure 2F and 2G). Immunohistochemistry staining showed significantly increased Ki-67 positive cells in tumors of *E. faecalis* and EF-CM treatment groups. Consistently, p-mTOR and its downstream phosphorylated proteins were also elevated (Figure 2H). Taken together, these results indicate that *E. faecalis* potentiates mTOR pathway activation to promote liver cancer progression.

### EVs of *E. faecalis* delivered through a dynamin-dependent endocytic process

Bacterial extracellular vesicles (EVs) are key mediators of bacteria-host interaction. Transmission electron microscopy (TEM) showed spherical EVs around *E. faecalis* (Figure 3A). Scanning electron microscopy and nanoparticle tracking analysis (NTA) revealed the isolated EVs as spherical bilayer membrane-enclosed vesicles with ∼160 nm diameter (Figure 3B), consistent with typical EVs features of Gram-positive bacteria^22^. 50 μg/mL EF-EVs were able to promote HCC cells proliferation (Figure 3C and 3D). EF-EVs could also promote HUVEC cells tube formation and migration (Figure 3E and 3F). Immunoblotting assays demonstrated that EF-EVs treatment increased the phosphorylation of mTOR and its downstream proteins P70S6K and 4EBP1 in HCC cells (Figure 3G). These findings suggest that EF-EVs are the potential effector in mediating mTOR activity in HCC.

**Figure 3.**
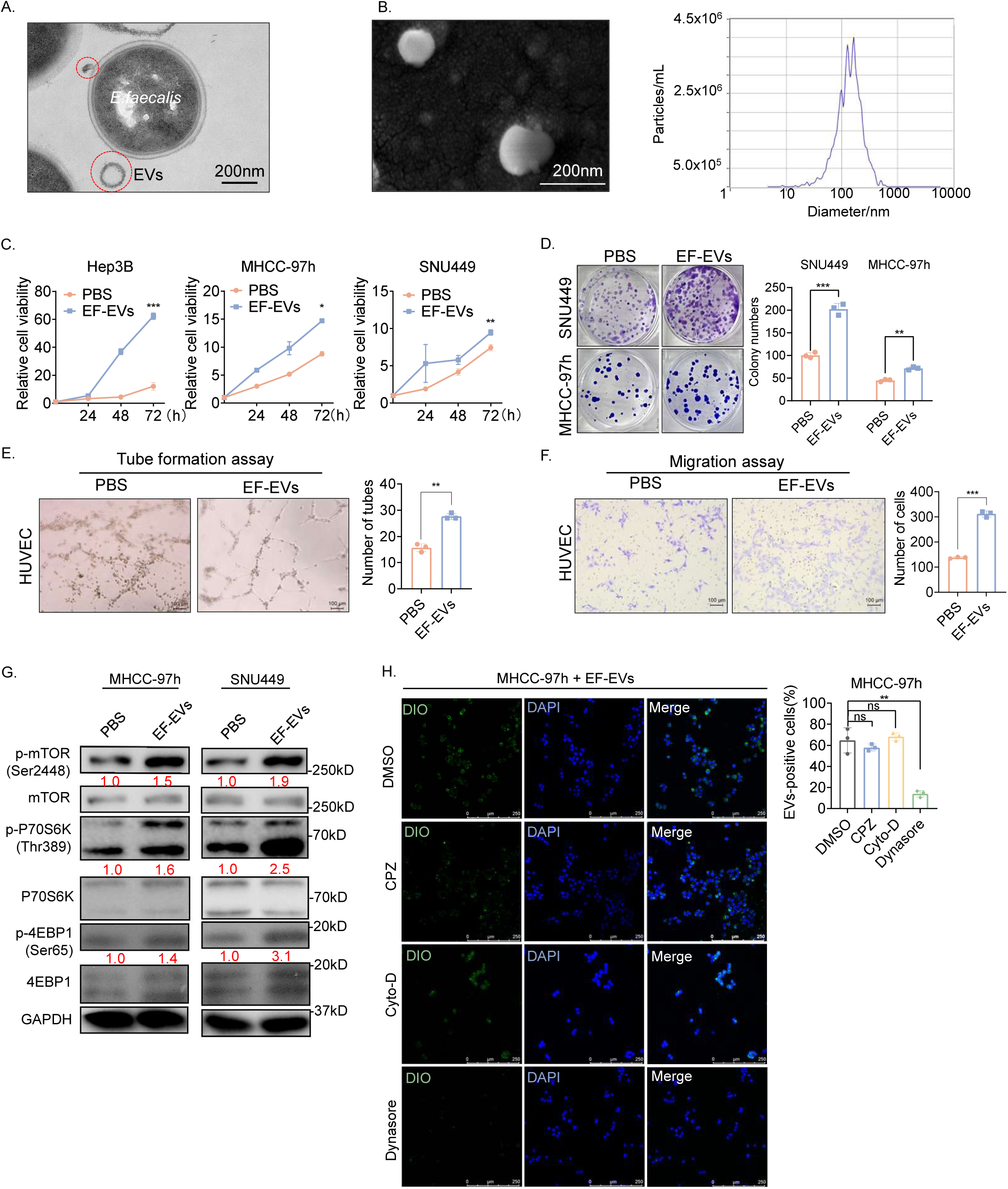
EVs of *E. faecalis* delivered through a dynamin-dependent endocytic process can promote liver cancer cell proliferation and activate mTOR. **(A)** Transmission electron microscopy (TEM) image of *E. faecalis* (cross-section) with secreted EVs (red circle). Scale bar = 200 nm. **(B)** Representative TEM image and nanoparticle tracking analysis (NTA) of EF-EVs. Scale bar = 400 nm. **(C&D)** CCK8 assay and Colony formation assay of HCC cells. **(E&F)** Tube formation and migration assay of HUVEC cells. Scale bar = 100 μm. **(G)** Protein levels of phosphorylation of mTOR, P70S6K and 4EBP1 in HCC cells under indicated treatments. **(H)** Confocal micrographs of MHCC-97h cells pretreated with indicated inhibitors (Chlorpromazine 5 µM, Cytochalasin D 2.5 µM, Dynasore 40 µM) for 1 h, followed by incubation with 20 μg/mL DiO-labeled EVs for 120 min. Quantification of EVs-positive cell were shown. Scale bars = 250 μm. Unpaired two-tailed Student’s *t* test was used in (C), (D), (E), (F) and (J). EF-EVs, *E. faecalis* extracellular vesicles; DIO, 3,3’-dioctadecyloxacarbocyanine perchlorates; DMSO, Dimethyl sulfoxide; CPZ, Chlorpromazine; CytoD, Cytochalasin D.

We labeled EF-EVs with lipophilic membrane dye 3,3’-dioctadecylindocarbocyanine perchlorate (DiO) (emits green fluorescence when entering cell membrane^23^) and treated MHCC-97h cells. To identify the specific endocytic pathway, we used inhibitors targeting different pathways: Chlorpromazine (clathrin-mediated), Dynasore (dynamin-mediated), and Cytochalasin D (actin-mediated). Immunofluorescence showed Dynasore significantly inhibited EVs endocytosis, while other inhibitors had no noticeable effect (Figure 3H). Thus, EF-EVs enter HCC cells via a dynamin-dependent endocytic process to exert their effects.

### EF-EVs promotes hepatocarcinogenesis in mice via enhancing mTOR signals

To validate the *in vivo* effects of EF-EVs, we established an orthotopic liver cancer model^24^ in C57BL/6 mice. After 2-week antibiotic pretreatment, mice were administered EF-EVs or PBS every other day (Figure 4A). For *in vivo* tracing, fluorescence images 90 minutes post oral gavage showed Cy7-prelabeled EF-EVs in mouse livers (Figure 4B). EF-EVs promoted HCC progression, as evidenced by stronger liver bioluminescence signals, larger orthotopic tumor volumes (Figure 4C-4F) and elevated blood biochemical markers ALT, AST, ALP (Figure 4G). Immunohistochemical staining showed EF-EVs gavage increased p-mTOR, p-P70S6K, p-4EBP1, HIF1α, CD31 levels and Ki-67 signal in tumor tissues (Figure 4H). These results indicated EF-EVs could promote HCC progression by enhancing the mTOR pathway.

**Figure 4.**
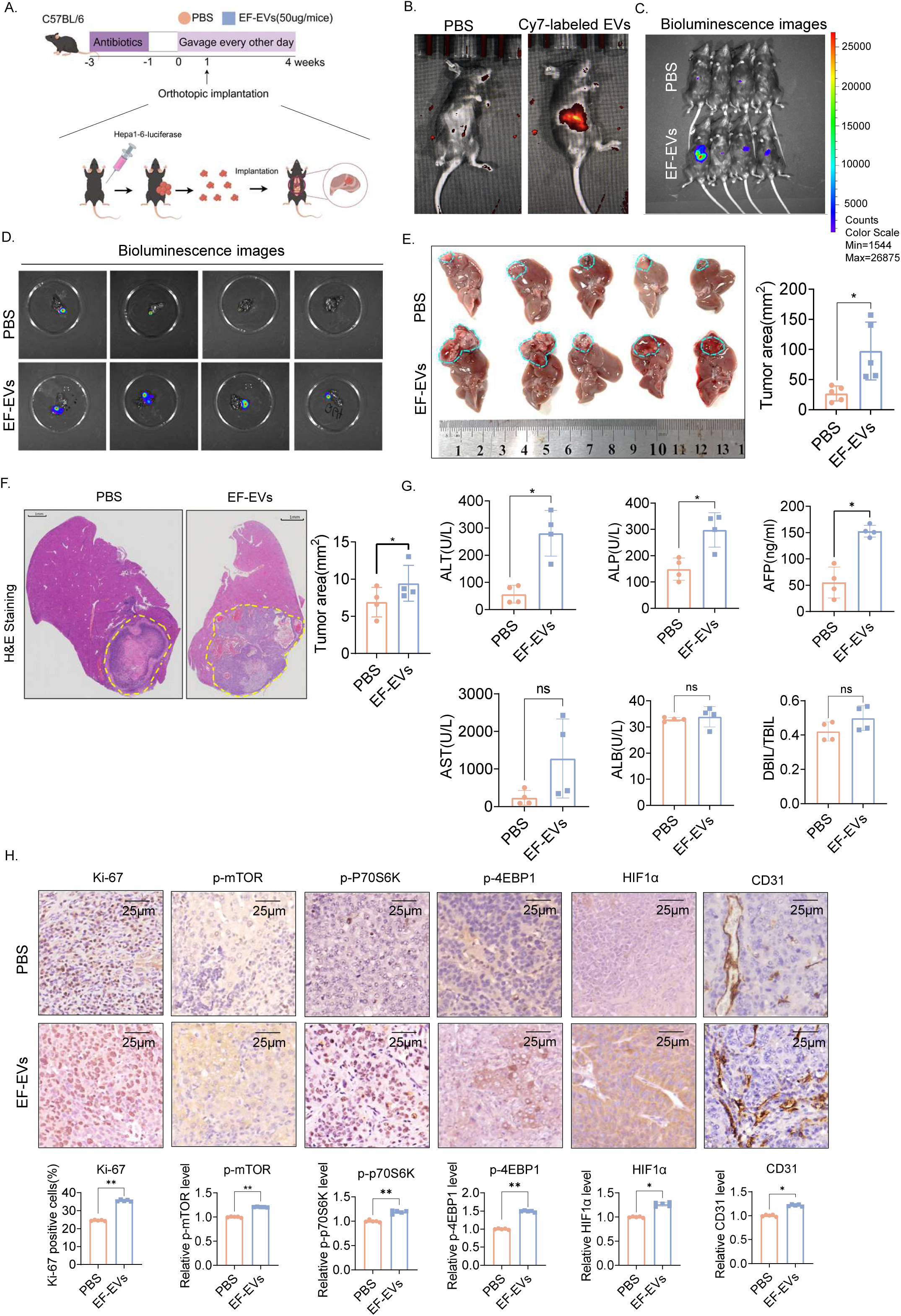
EF-EVs promotes hepatocarcinogenesis in mice and causes pathological conditions. **(A)** Scheme of EF-EVs mediated liver tumorigenesis experiment. **(B)** Representative fluorescence images of mice 90 min after gavage with Cy7-labeled EF-EVs (right) or PBS (left). **(C&D)** Bioluminescence images of mouse whole body and isolated livers at the end of the experiments. **(E)** Images of livers harvested from EF*-*EVs and PBS (n = 5) group. The blue dashed lines indicate the tumor borders. **(F)** Representative images of H&E staining on liver tissue sections (left). **(G)** Quantification of serum biomarkers levels from two groups. **(H)** Representative images of immunohistochemical staining in tumor samples. Scale bar = 25 μm. Unpaired two-tailed Student’ *t* test was used in (E), (F), (G) and (H). ALT, Alanine Transaminase; ALP, Alkaline phosphatase; AFP, α-fetoprotein; AST, Aspartate Aminotransferase; ALB, albumin; DBIL, direct bilirubin; TBIL, total bilirubin.

16S rRNA profiling analysis showed that EF-EVs led to a significant reduction in the overall abundance of the gut microbiota (Figure S2A), together with altered gut microbiota composition (Figure S2B, S2C and S2D). Notably, it reduced colonization of probiotics with known gut health benefits, while promoting colonization of *E. faecalis* itself and harmful bacteria (Figure S2E and S2F). These findings suggest EF-EVs gavage may exert complex effects on gut microbiota, potentially disrupting beneficial-harmful bacteria balance and altering the overall gut ecosystem.

### Obg derived from *E. faecalis* EVs is a GTPase physically interacts with mTOR

Bacteria-derived proteins, DNA, and RNA can interact with host cell molecules and modulate key signaling pathways, influencing tumor progression^25, 26^. Pretreating EF-EVs with universal nuclease (deplete DNA/RNA) or heat-killed (denature proteins) showed heat-killed EVs lost HCC-promoting activity, while nuclease-treated EVs remained active (Figure 5A), suggesting that EF-EVs exert oncogenic effects via proteins. Mass spectrometry analysis of EF-EVs protein components revealed enrichment of ribosome-related proteins involved in protein synthesis (Figure S3A and S3B). Furthermore, mTOR co-IP mass-spectrometry identified 547 EF-EVs-derived proteins/peptides (Figure 5B). Hypothesizing an EF-EVs GTPase might mimic Ras-family GTPase Rheb (which activates lysosomal Mtorc1) in promoting HCC, several EF-EVs GTPases were identified, with spo0B-associated GTP-binding protein (Obg) being particularly notable. Obg proteins (TRAFAC class of P-loop GTPases) are involved in DNA replication, ribosome maturation, and stress adaptation^27^. EF-Obg has an N-terminal domain, a non-conserved C-terminal region, and a central G domain common to all Obg proteins^27^ (Figure 5C). Obg proteins possess a Ras-like folds G domain with five conserved motifs for GDP and GTP recognition and hydrolysis, similar to Ras or Rheb^28^. Structural modeling predicted EF-Obg forms an mTOR complex analogous to Rheb (Figure 5D). Exogenous expression via the transcription and translation (TNT) system confirmed direct EF-Obg-mTOR binding (Figure 5E), further validated in HEK293T and MHCC-97h cells (Figure 5F). Proximity ligation assays demonstrated in vivo binding (Figure 5G). Collectively, these data indicate direct interaction between mTOR and EVs-derived EF-Obg.

**Figure 5.**
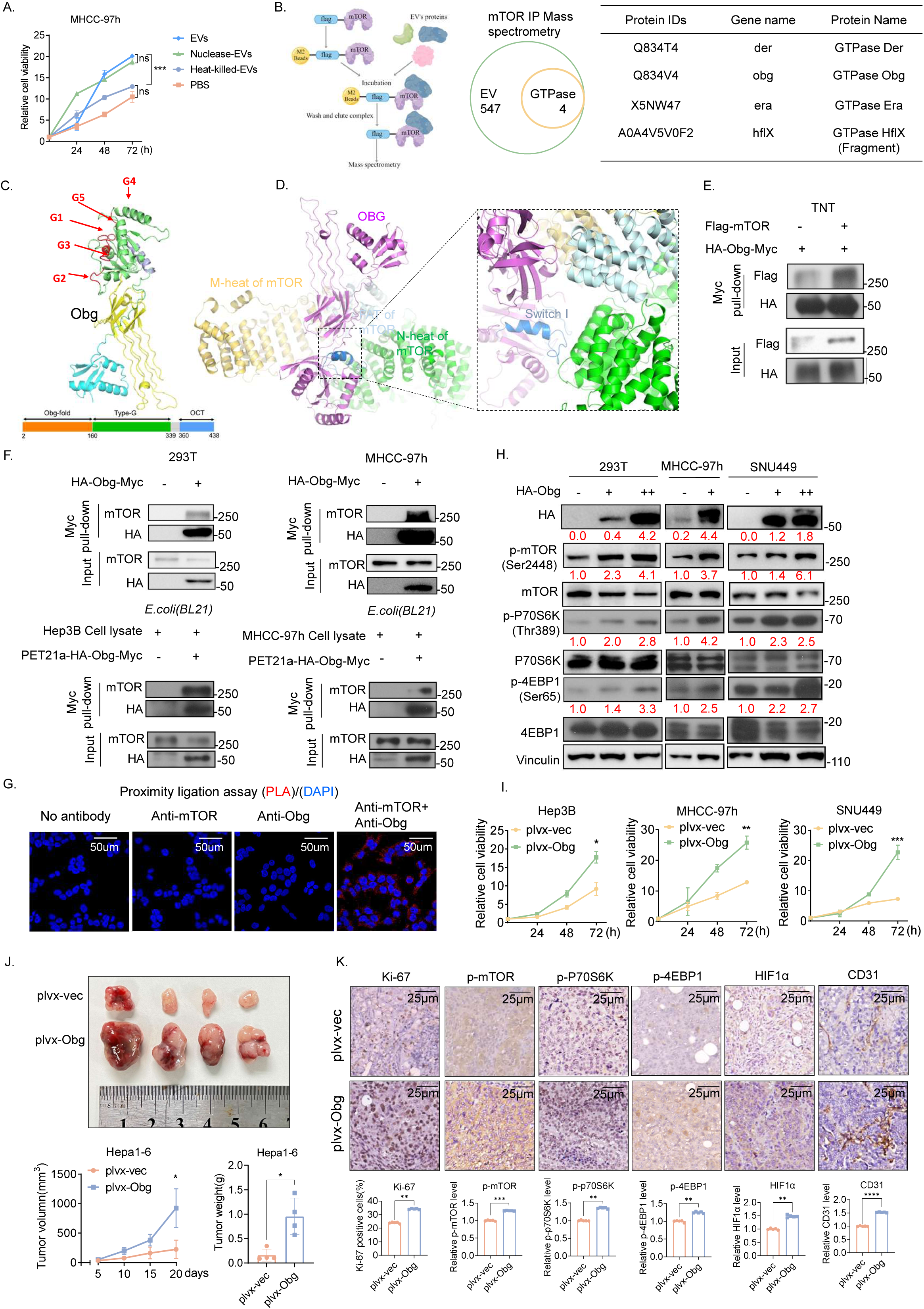
EF-Obg derived from EF-EVs is a GTPase physically interacts with mTOR. **(A)** The growth curves of MHCC97h cells with indicated treatments were measured by CCK8 assay. **(B)** Schematic diagram of mTOR IP Mass spectrometry. Four GTPases were identified (right). **(C&D)** GTPase-Obg protein structure with domains (C) and Obg-mTOR interaction structure (D) visualized by PyMOL. The red arrows showed the GTP binding sites (G1-G5). **(E)** *In vitro* co-IP assays were performed using protein products from transcription and translation (TNT) assays. **(F)** Co-IP assays were performed using exogenously expressed HA-Obg-Myc in 293T/MHCC-97H cells, or using bacterially expressed PET-21a-HA-Obg-Myc mixed with Hep3B/MHCC-97H cell lysates. **(G)** Representative image results of proximity ligation assay (PLA) using antibodies against mTOR and Obg. The red signals indicate interactions between mTOR and Obg. Scale bar = 50 μm. **(H)** Protein levels of phosphorylation of mTOR, P70S6K, and 4EBP1 after HA-Obg overexpression in indicated cells. **(I)** Cell growth curves of indicated HCC cells overexpressing Obg. Cell growth was measured by CCK8 assay. **(J)** C57BL/6 female mice were subcutaneously injected with 5×10^5^ Hepa1-6 cells overexpressing Obg (plvx-Obg) or control (plvx-vec). Representative tumor picture, tumor volumes, and tumor weights were shown (n = 4). **(K)** Representative images of immunohistochemical staining from tumor samples. Scale bar = 25 μm. One way ANOVA test was used in (A). Unpaired two-tailed Student’s *t* test was used in (I), (J) and (K).

### *E. faecalis obg* gene is critical for *E. faecalis* to activate mTOR to promote hepatocarcinogenesis

EF-Obg binding dose-dependently enhanced mTOR pathway activation, confirming its role in mTOR regulation (Figure 5H). Consistently, EF-Obg overexpression promoted HCC cell proliferation (Figure 5I). In a mouse model, subcutaneous injection of Hepa1-6 cells stably overexpressing EF-Obg into C57BL/6 mice showed increased tumor growth and weight (Figure 5J), with IHC staining confirming EF-Obg-mediated mTOR pathway activation (Figure 5K). These results indicate that EF-Obg activates mTOR pathway to promote liver tumorigenesis. PyMOL comparison with classic mTOR-activating GTPase Rheb revealed EF-Obg has a conserved G1 site amino acid sequence (Figure 6A). A G1 site missense mutation generated the Obg^8A^ mutant (Figure 6B). Co-IP experiments showed the Obg^8A^ mutant lost mTOR-binding ability (Figure 6C). Consistently, the Obg^8A^ mutant failed to activate mTOR, whereas wildtype (WT) Obg retained mTOR-activating activity (Figure 6D).

**Figure 6.**
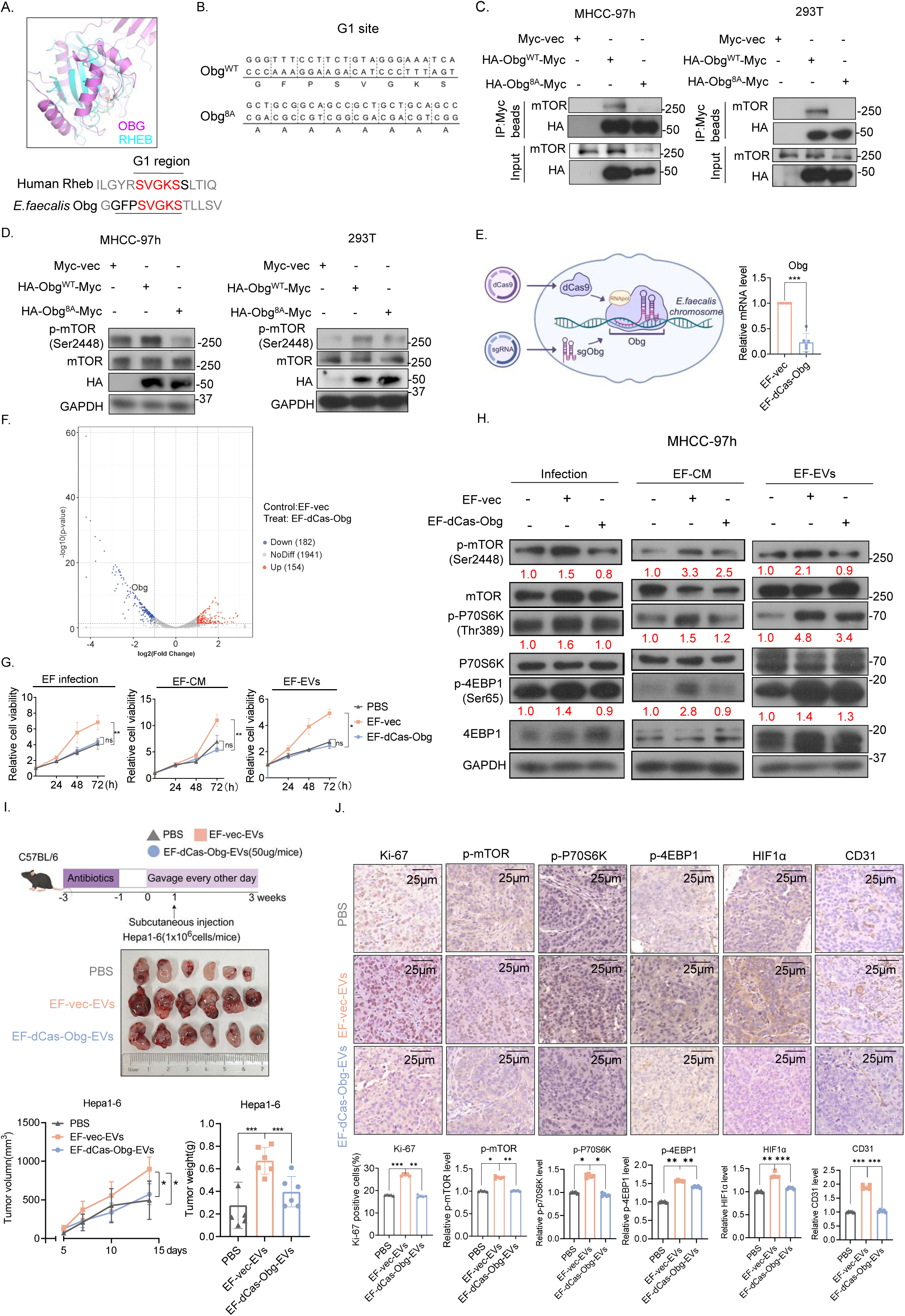
*E. faecalis obg* gene is critical for *E. faecalis* to activate mTOR to promote hepatocarcinogenesis. **(A)** Comparison of GTP binding sites 1 (G1) homology between Obg and Rheb. Aligned amino acid sequences in G1 were highlighted. **(B)** The detailed nucleotide sequence and the translated amino acids of the G1 site from EF-Obg wild-type (Obg^WT^) and G1 site mutant (Obg^8A^). **(C)** The interaction between mTOR and Obg^WT^ or Obg^8A^ was determined by co-IP assay. **(D)** The phosphorylation levels of mTOR following Obg^WT^ or Obg^8A^ overexpression were shown. **(E)** Schematic diagram illustrating the knockdown of Obg by CRISPR interference system. Obg mRNA level was measured by qPCR. **(F)** Volcano blot showed differentially expressed genes in EF-dCas-Obg strain compared with EF-vec strain, as revealed by RNA-sequencing analysis. **(G)** The growth curves of MHCC-97h cells with indicated treatments were measured by CCK8 assays. **(H)** Protein levels of phosphorylation of mTOR, P70S6K, 4EBP1 in HCC cells with indicated treatments. **(I)** Schematic diagram of animal experiments. Representative tumor image, tumor volumes, and tumor weights were shown (n = 6). **(J)** Representative images of immunohistochemical staining from tumor samples of indicated group. Scale bar = 25 μm. Unpaired two-tailed Student’s *t* test was used in (E); One-way ANOVA was used in (G), (I) and (J). G1 site, GTP-binding site 1.

To verify the critical role of the *E. faecalis obg* gene, we used the CRISPR interference system^21^ to engineer an *E. faecalis obg* knockdown strain (EF-dCas-Obg) (Figure 6E). Target EF-Obg sequences for sgRNA predicted by CHOPCHOP are shown in Table S4. qPCR and RNA sequencing validated successful *obg* knockdown (Figure 6E and 6F). Notably, EF-dCas-Obg exhibited retarded growth, suggesting *obg* is essential for *E. faecalis* (Figure S4A). NTA analysis revealed decreased concentrations of secreted EVs particles and EVs protein content in EF-dCas-Obg compared with the EF-vec strain, while EVs size remained unchanged (Figure S4B and 4C). GO and KEGG analyses indicated that *obg* knockdown affects biological functions such as transmembrane transport and fatty acid synthetase activity (Figure S4D and S4E).

EF-dCas9-Obg strain had reduced ability to promote HCC cell proliferation compared with the EF-vec strain (Figure 6G). Consistently, EF-dCas9-Obg strain had compromised activating effect on the mTOR signaling pathway when compared to EF-vec strain (Figure 6H). Similarly, EVs from EF-dCas9-Obg had a diminished effect on promoting liver cancer growth in mice compared with EF-vec (Figure 6I). IHC staining showed that EVs from EF-dCas9-Obg lost their ability to activate the mTOR pathway (Figure 6J). 16S sequencing revealed that the EF-dCas9-Obg strain altered the composition of the mouse gut microbiota (Figure S5A-S5C). Notably, it increased the abundance of *Akkermansia*, a well-known probiotic for liver disease (Figure S5D and S5E), indicating that *obg* knockdown changes the mouse microbial landscape. Collectively, these results establish a causal link between microbiota dysbiosis and liver carcinogenesis, demonstrating that *obg* gene knockdown mitigates *E. faecalis*’ promoting role in activating mTOR and promoting hepatocarcinogenesis.

### EF-Obg expression level in liver cancer tissue correlates with poor prognosis in HCC patients

To explore the clinical significance of our findings, we investigated the expression level of EF-Obg in liver tumor tissues and corresponding normal tissues. qPCR has demonstrated the high level of *obg* gene expression in HCC (Figure 7A). A monoclonal antibody against EF-Obg GMVAFRREKYVPD sequence was generated using hybridoma technology (Figure S6A-S6D). The sensitivity and specificity of the monoclonal antibody were validated on tissue sections (Figures 2H and 4H) from previous animal experiment (Figure S6E). Notably, higher levels of EF-Obg were observed in tumor tissues when compared with normal tissues, concurrent with elevated levels of p-mTOR (Figures 7B and C). IHC staining using this monoclonal antibody against EF-Obg in a panel of 100 HCC and normal tissue specimens demonstrated that patients with high EF-Obg expression had significantly reduced overall survival and recurrence-free survival compared to those patients with low Obg expression (Figure 7D and 7E). Clinical association study demonstrates that EF-Obg expression was significantly correlated with HCC tumor number, tumor size, TNM (Tumor, Node, Metastasis) stage and BCLC (Barcelona Clinic Liver Cancer) stage (Supplemental Table 1).

**Figure 7.**
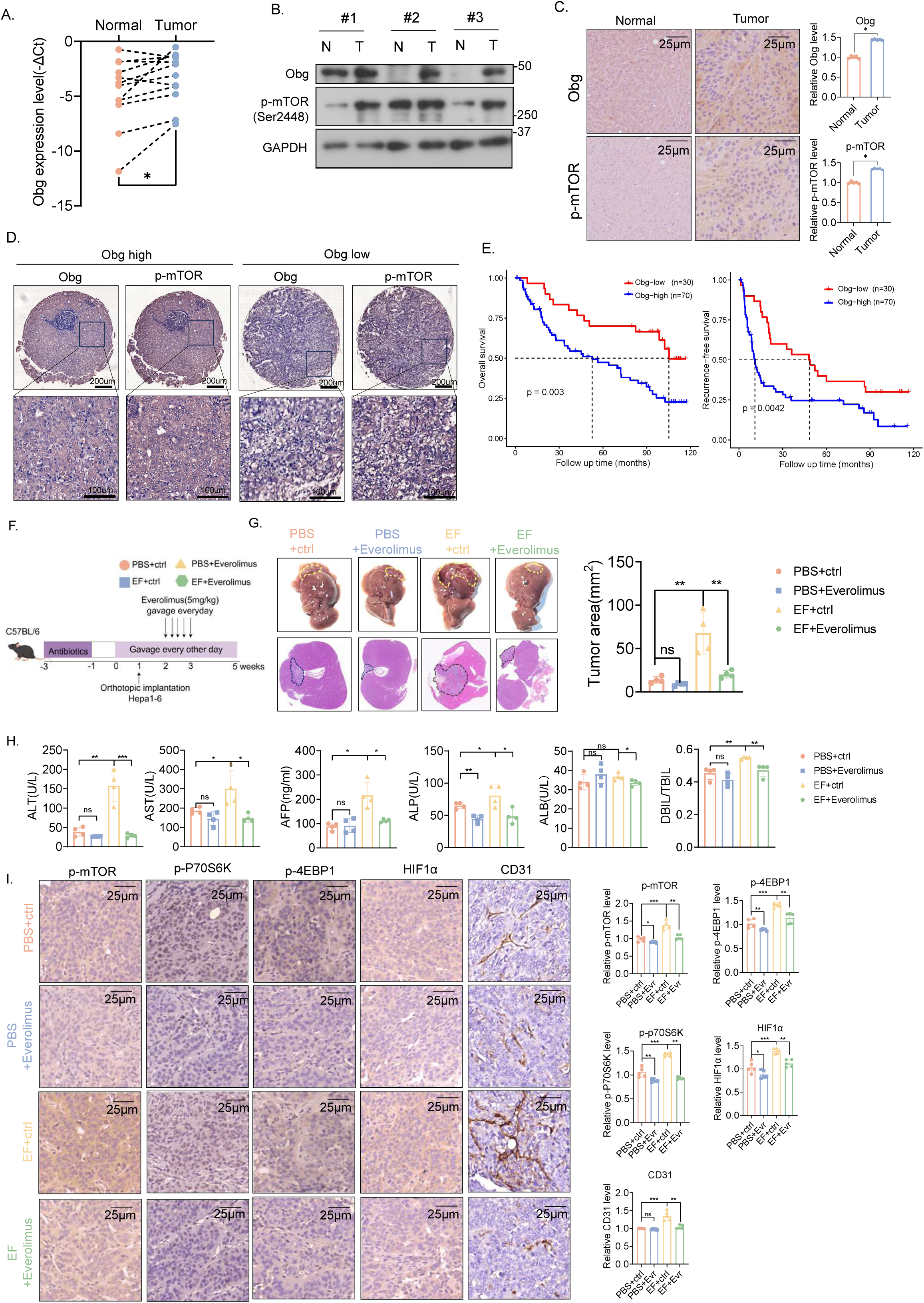
High EF-Obg expression level in liver cancer tissue correlates with poor survival, and can be targeted for therapy using mTOR inhibitor Everolimus. **(A)** Relative expression of EF-Obg in tissues based on qPCR. **(B)** Protein levels of EF-Obg and p-mTOR in paired liver tumor tissues and normal tissues. N, normal tissues; T, liver tumor tissues. **(C)** Representative images of immunohistochemical staining of EF-Obg (using anti-EF-Obg antibody) and p-mTOR in tissues. Scale bar = 25 μm. **(D)** Representative IHC staining images showing high and low expression of EF-Obg in human HCC and normal tissues from tissue microarray (TMA). The matched p-mTOR staining was shown on the right. **(E)** Kaplan‒Meier survival curves were generated to estimate the overall survival and recurrence-free survival differences between Obg-high and Obg-low groups using a log rank test. N = 100. **(F)** Everolimus treatment scheme of EF-mediated liver tumorigenesis. Everolimus or control was orally administered to the mice every day one week after the tumor implantation. **(G)** Representative images of the Everolimus-treated tumors under *E. faecalis* administration, H&E staining of the tumors (n = 4). **(H)** Serum levels of serum biomarkers of mice after indicated treatments. **(I)** Representative images of immunohistochemical staining. Scale bar = 25 μm. Paired *t*-test was used in (A). Unpaired two-tailed Student’s *t* test was used in (C). Log rank test was used to calculate the *p* value in (E). One-way ANOVA test was used in (G), (H) and (I).

### EF-mediated mTOR activation in liver cancer can be targeted for therapy using mTOR inhibitor Everolimus

Given that Everolimus, a rapalogue, is a potent inhibitor of mTOR kinase, we then investigated whether this drug has a good inhibitory effect on liver cancer colonized with *E. faecalis*. We established an orthotopic mouse liver cancer model (Hepa1-6) in C57BL/6 mice gavaged with *E. faecalis* to investigate the treatment efficacy of Everolimus (Figure 7F). The results showed that in the presence of *E. faecalis* colonization, Everolimus significantly suppressed tumor growth and liver damage, as evidenced by reduced tumor area and serum biomarkers (ALT and AST), while it showed no significant efficacy against liver cancer in the absence of *E. faecalis* colonization (Figure 7G and 7H). IHC staining confirmed the efficient effect of Everolimus in inhibiting mTOR pathway in EF-colonized mouse liver cancer model (Figure 7I). These data suggest that EF-Obg is an actionable biomarker, and indicate that Everolimus treatment inhibits EF-Obg-mediated mTOR signaling axis and may be considered as a therapeutic strategy for liver cancer patients with *E. faecalis* colonization.

## Discussion

Although increasing evidence links gut microbiota to HCC through the gut-liver axis^29, 30^, the direct impact and underlying mechanisms remains largely unknown. Here, we demonstrate that tumor-resident *E. faecalis* activates mTOR signaling pathway via secreting Obg GTPase containing extracellular vesicles. Our study sheds light on *E. faecalis* signaling communications with host cells and reveals how mTOR is activated by *E. faecalis* to promote HCC tumorigenesis, focusing on a microbial mTOR regulator.

### *E. faecalis* acts as an oncobacterium of liver cancer via activation of mTOR

*E. faecalis* was found to be highly enriched in the feces of patients with liver diseases. In addition to its colonization in the gut, *E. faecalis* is able to colonize several cancer tissues^3, 15, 31^; Our findings revealed that the abundance of *E. faecalis* in HCC tissues is verified and clinically significant, suggesting its potential as a biomarker for HCC diagnosis and prognosis (Figure 1). It is possible that other non-characterized bacteria species in our study may also play roles in promoting HCC and can serve as tumor marker, but they remain to be further explored. Our results demonstrate that EF-derived EVs have oncogenic impact in the development of liver cancer, particularly providing insight into the role of EF-EVs in regulation of host mTOR pathway. Further investigation into the other potential pathways regulated by *E. faecalis*, such as RelA and STK33 regulations implied in our studies (Figure2), in HCC tumorigenesis could be critical and need to be further determined as interventions based on intratumoral microbiota regulations will be of great potential to rational cancer therapy.

### EVs-derived EF-Obg is an inter-kingdom activator of mTOR

EVs facilitate host-bacteria communication by delivering proteins or nucleosides^23, 25, 32, 33^. We show EF-EVs act as key biological shuttles for inter-kingdom communication, transferring bacterial signals to host. Our study highlighted EF-derived EVs carry crucial molecules implicated in mTOR signaling, notably EF-Obg GTPase that regulates mTOR. EF-Obg proteins have conserved Ras-like G domain (G1) motifs, which mediate guanine nucleotide binding and hydrolysis process – critical for bacterial viability (e.g. *C. crescentus*) and bacterial persistence under nutrient starvation^34^, similar to mTOR’s nutrition regulatory role. Modeling studies demonstrate that EF-Obg binds mTOR just like Rheb. Usually, Obg proteins demonstrate a high guanine nucleotide exchange rate (10^3^- to 10^5^-fold faster than that of eukaryotic Ras-like GTPases, possibly faster than Rheb^35^), suggesting EF-Obg may activate mTOR more efficiently than host Rheb, accelerating HCC growth.

In a mouse liver cancer model, *E. faecalis* with *obg* gene knockdown causes attenuated mTOR activity, recapitulating EF-Obg’s positive impact on mTOR. Interestingly, human Obg homolog protein human Obg-like ATPase1(hOLA1) is overexpressed in various cancers^36, 37^, regulating protein synthesis, integrated stress response (ISR) and acting as a novel GSK3β inhibitor^38^. Whether hOLA1 regulates mTOR or interacts with EF-Obg in oncogenesis remains unclear.

### *E. faecalis* Obg is a tumor marker, and mTOR inhibition treatment can be a rational cancer therapy for HCC colonized with tumor-resident *E. faecalis*

Hepatocellular carcinoma (HCC) results from various accumulation of multiple genomic and epigenomic alterations, leading to deregulation of signaling pathways, including AKT/mTOR signaling^39^. The PTEN/PI3K/Akt/mTOR axis and downstream target ATP-binding cassette subfamily C member 4 (Abcc4) are involved in hepatocarcinogenesis^40^, but mTOR activation in HCC is not fully characterized. Importantly, we produced EF-Obg antibody and detected Obg expression in liver tumors. Our results show that EF-Obg expression levels correlate with mTOR activation and poor overall survival, suggesting they could be parallel prognostic tumor markers, with EF-Obg antibodies serving as a diagnostic tool. Since mTOR inhibition significantly dampens HCC growth and improves survival, mTOR inhibitors may improve HCC treatment^41^. Our study explored the therapeutic potential of mTOR kinase inhibitor Everolimus in EF-colonized liver cancer models. Since EF-Obg regulates mTOR to potentiate cancer protein translation and angiogenesis, Everolimus might lead to a better efficacy in treating EF-colonized liver cancer. Indeed, Everolimus has excellent efficacy in inhibiting mTOR activity caused by the *E. faecalis* in liver cancer. Thus, these studies bear important therapeutic implications for HCC with tumor-resident *E. faecalis*. Future screening of inhibitors targeting EF*-*Obg-mTOR interaction (e.g. competing peptides) could offer another treatment strategy.

In conclusion, our results uncover an unconventional pathway that bacterial protein activates mTOR to promote tumor growth. The effect of EF-Obg in enhancing mTOR activation via serving as a GTPase illustrates a cross-kingdom regulation regarding the mTOR dysregulation caused by microbiota dysbiosis during HCC progression. Treatment strategies that block mTOR activation by Everolimus can be further employed as a rational cancer therapy for HCC colonized with tumor-resident *E. faecalis*.

## Authors Contributions

Mong-Hong Lee and Xiangqi Meng conceived and designed the research. Ning Ma, Xiaoshan Xie and Jiarui Wang performed most of the biochemical and molecular experiments, with the assistance from Xiaoling Huang, Yue Wei, Qihao Pan, Boyu Zhang, Jiaying Zheng, Peng Zhang. Zhikai Zheng and Zhongguo Zhou ascertained and processed clinical specimens. Zhikai Zheng performed the bioinformatics analysis. Ning Ma and Haidan Luo constructed the obg knockdown strain by CRISPR interference under Xue Liu’s guidance. Zhi-Min Zhang predicted Obg-mTOR binding/interactions domain. Ning Ma, Jiarui Wang Xiaoshan Xie, Huilin Jin and Xijie Chen performed mice experiments. Ning Ma and Jiarui Wang conducted the bioinformatics analyses. Xijie Chen, Xiaoling Huang, Qihao Pan, Yue Wei, Boyu Zhang, Jiaying Zheng and Fenghai Yu contributed to discussion and data interpretation. Ning Ma, Xiangqi Meng, and Mong-Hong Lee wrote the manuscript. All authors have read and approved the final manuscript.

## Data avaliability Statement

Protocol details, further information and requests for resources and reagents can be directed by the lead contact Prof Mong-Hong Lee. The 16S data, transcriptome data and mass spectrometry proteomics data will be further uploaded. This paper does not report original code.

## Impact and Implications

While prior studies have consistently reported elevated abundance of *E. faecalis* in patients with HCC and suggested its pro-tumorigenic role, our results uncover an unconventional pathway that bacterial protein activates mTOR to promote tumor growth. The effect of EF-Obg in enhancing mTOR activation via serving as a GTPase illustrates a cross-kingdom regulation regarding the mTOR dysregulation caused by microbiota dysbiosis during HCC progression. Treatment strategies that block mTOR activation by Everolimus can be further employed as a rational cancer therapy for HCC colonized with tumor-resident *E. faecalis*.

