## supplemental figures and tables for "*Enterococcus faecalis* delivers Obg GTPase via extracellular vesicles to instigate mTOR activity and promote HCC tumorigenesis": supplemental files-20250714.pdf

Figure S1

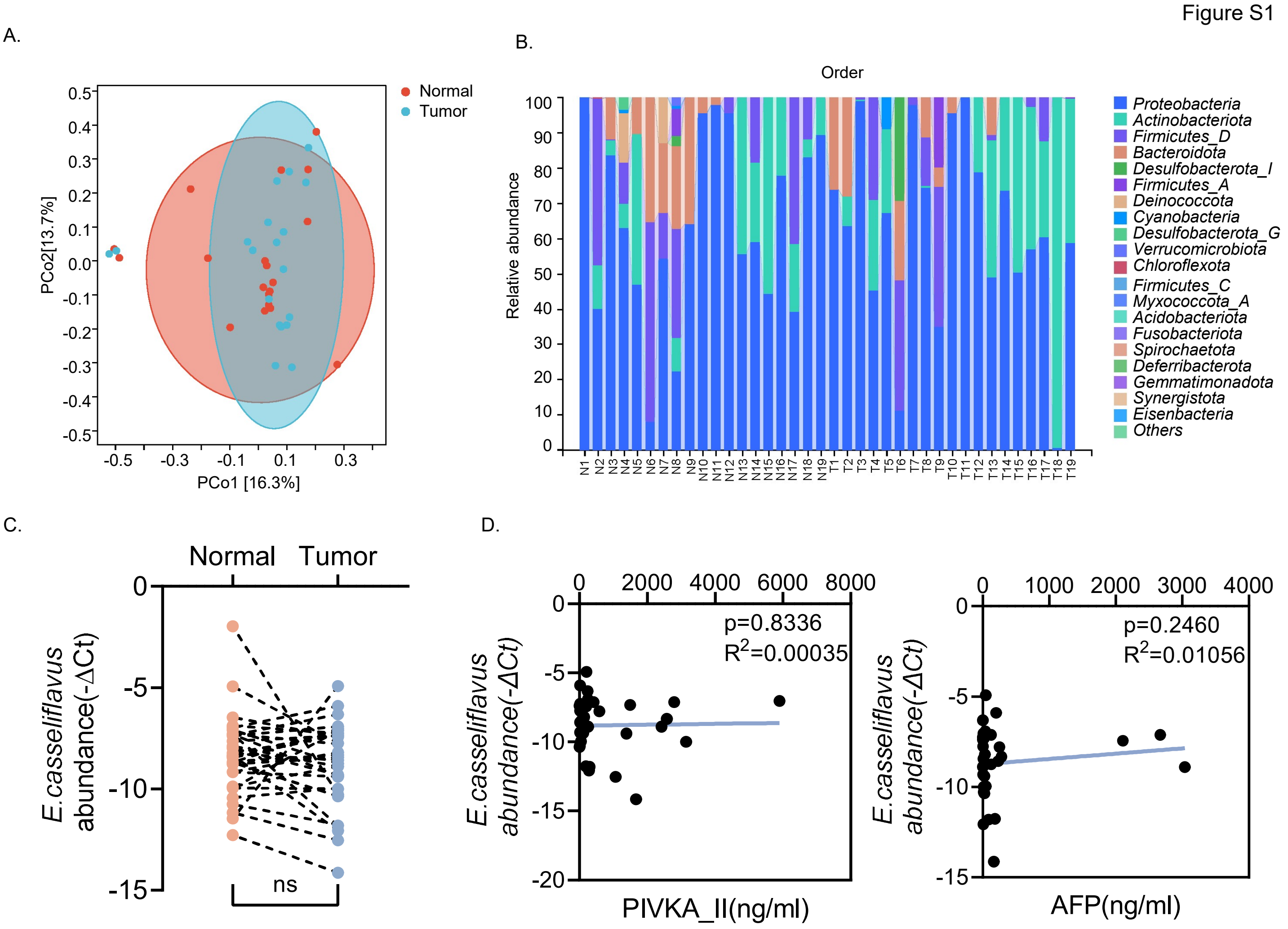

**Supplemental Figure 1 The composition of tissue-resident bacteria in liver tumor and normal tissues.**

(A) Principal Coordinate Analysis (PCoA) based on unweighted UniFrac analysis of extracted DNA from tissue-resident bacteria in tumor and normal tissues. Each dot represents a single sample.

(B) Relative abundance of tissue-resident microbiota in tumor and normal tissues at Order level based on 16S sequencing.

(C) Relative abundance of *E.casseliflavus* in liver tumor tissues and normal tissues.

(D) Spearman's correlation between -ΔCt values of *E.casseliflavus* with serum PIVKA_II level and AFP level in HCC patients.

ns, not significant. Paired *t*-test was used in (C).

Figure S2

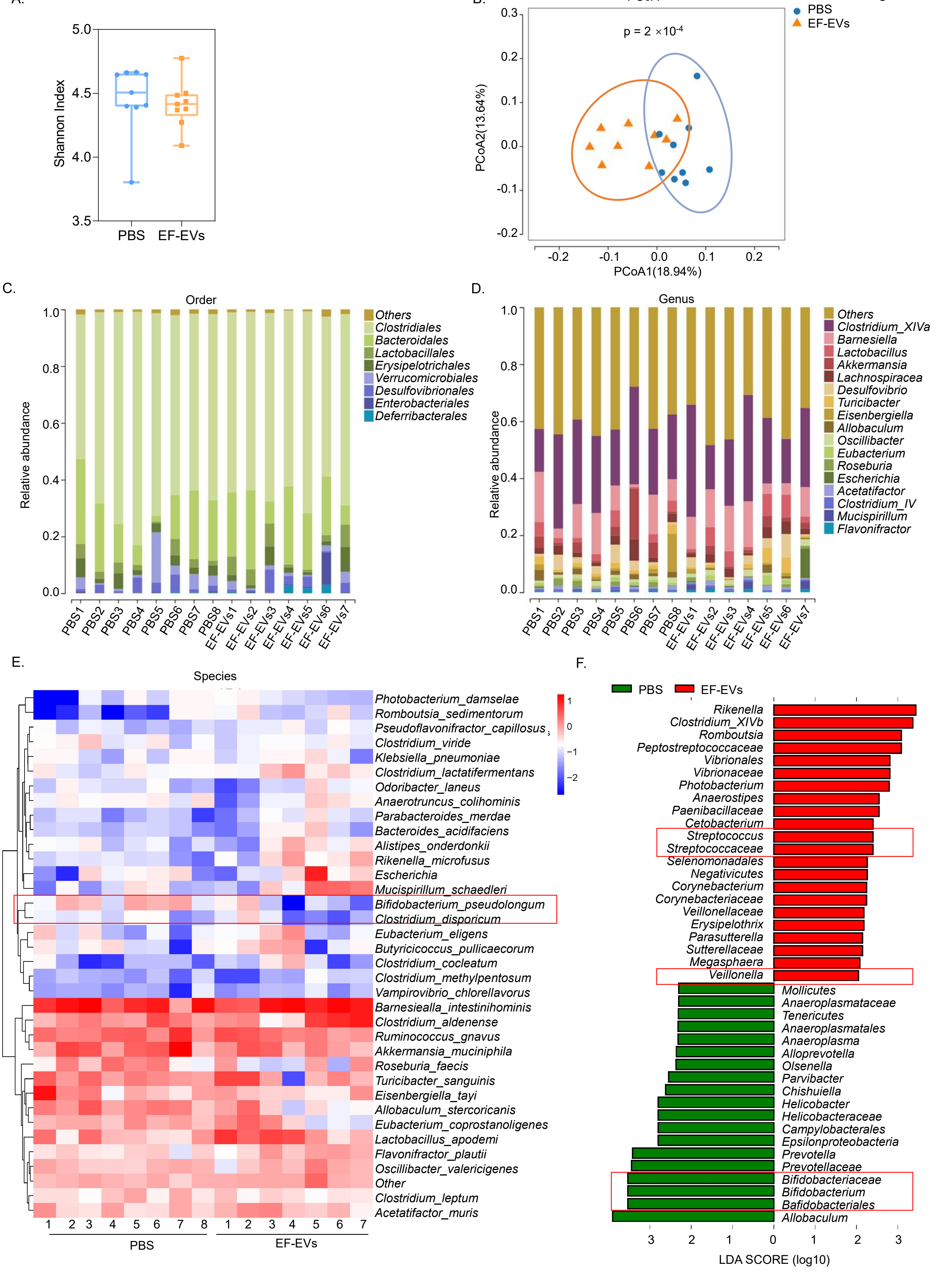
A.
Shannon Index
PBS
EF-EVs
B.
PCoA
p = 2 ×10-4
PBS
EF-EVs
PCoA1(18.94%)
PCoA2(13.64%)
C.
Order
Relative abundance
Others
Clostridiales
Bacteroidales
Lactobacillales
Erysipelotrichales
Verrucomicrobiales
Desulfovibrionales
Enterobacteriales
Deferribacterales
PBS1 PBS2 PBS3 PBS4 PBS5 PBS6 PBS7 PBS8 EF-EVs1 EF-EVs2 EF-EVs3 EF-EVs4 EF-EVs5 EF-EVs6 EF-EVs7
D.
Genus
Relative abundance
Others
Clostridium_XIVa
Barnesiella
Lactobacillus
Akkermansia
Lachnospiracea
Desulfovibrio
Turicibacter
Eisenbergiella
Allobaculum
Oscillibacter
Eubacterium
Roseburia
Escherichia
Acetatifactor
Clostridium_IV
Mucispirillum
Flavonifractor
E.
Species
Photobacterium_damselae
Romboutsia_sedimentorum
Pseudoflavonifractor_capillosus
Clostridium_viride
Klebsiella_pneumoniae
Clostridium_lactatifermentans
Odoribacter_laneus
Anaerotruncus_colihominis
Parabacteroides_merdae
Bacteroides_acidifaciens
Alistipes_onderdonkii
Rikenella_microfusus
Escherichia
Mucispirillum_schaedleri
Bifidobacterium_pseudolongum
Clostridium_disporicum
Eubacterium_eligens
Butyricicoccus_pullicaecorum
Clostridium_cocleatum
Clostridium_methylpentosum
Vampirovibrio_chlorellavorus
Barnesiealla_intestinihominis
Clostridium_aldenense
Ruminococcus_gnavus
Akkermansia_muciniphila
Roseburia_faecis
Turicibacter_sanguinis
Eisenbergiella_tayi
Allobaculum_stercoricanis
Eubacterium_coprostanoligenes
Lactobacillus_apodemi
Flavonifractor_plautii
Oscillibacter_valericigenes
Other
Clostridium_leptum
Acetatifactor_muris
PBS
EF-EVs
F.
PBS
EF-EVs
Rikenella
Clostridium_XIVb
Romboutsia
Peptostreptococcaceae
Vibrionales
Vibrionaceae
Photobacterium
Anaerostipes
Paenibacillaceae
Cetobacterium
Streptococcus
Streptococcaceae
Selenomonadales
Negativicutes
Corynebacterium
Corynebacteriaceae
Veillonellaceae
Erysipelothrix
Parasutterella
Sutterellaceae
Megasphaera
Veillonella
Mollicutes
Anaeroplasmataceae
Tenericutes
Anaeroplasmatales
Anaeroplasma
Alloprevotella
Olsenella
Parvibacter
Chishuiella
Helicobacter
Helicobacteraceae
Campylobacterales
Epsilonproteobacteria
Prevotella
Prevotellaceae
Bifidobacteriaceae
Bifidobacterium
Bafidobacteriales
Allobaculum
LDA SCORE (log10)

**Supplemental Figure 2 *E. faecalis* EVs change the intestinal microecological environment of mice.**

(A) 16S sequencing of mice feces from EF-EVs or PBS treatment group. Shannon index showed the alpha diversity of microbiota between 2 groups.

(B) PCoA based on un-weighted UniFrac analysis of mice feces microbiota in two groups. Each dot represents a single sample.

(C) Relative abundance of fecal microbiota in samples from EF-EVs or PBS treatment group at Order level.

(D) Relative abundance of fecal microbiota in samples from EF-EVs or PBS treatment group at Genus level.

(E) Heatmap showing fecal bacteria species distribution in samples from EF-EVs or PBS treatment group. The red box highlights species that were significantly decreased in the EF-EVs group.

(F) Linear discriminant analysis (LDA) statistical method was used to identify the most differentially abundant taxa between two groups (LDA > 2 and $p < 0.05$). The red box highlights altered species that have been reported to be associated with HCC.

Wilcoxon test was used to calculate the *p* value in (A). Unpaired two-tailed Student's *t* test was used in (B).

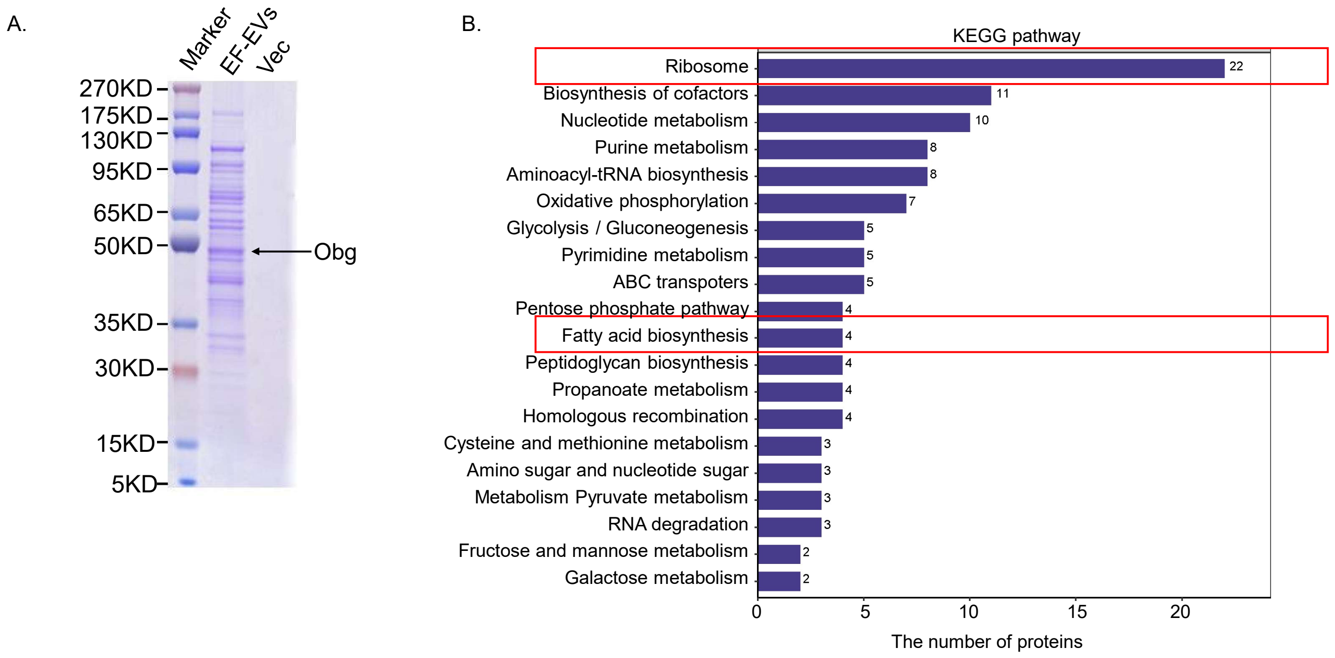

**Supplemental Figure 3 Proteomic analysis of extracellular vesicles of *E. faecalis***

(A) Coomassie Blue staining of EF-EVs protein bands based on SDS-PAGE.

(B) The Kyoto Encyclopedia of Genes and Genomes (KEGG) annotation results of the identified proteins from EF-EVs. The red boxes highlight the most abundant proteins belonging to the ribosome group and fatty acid biosynthesis group.

Figure S4

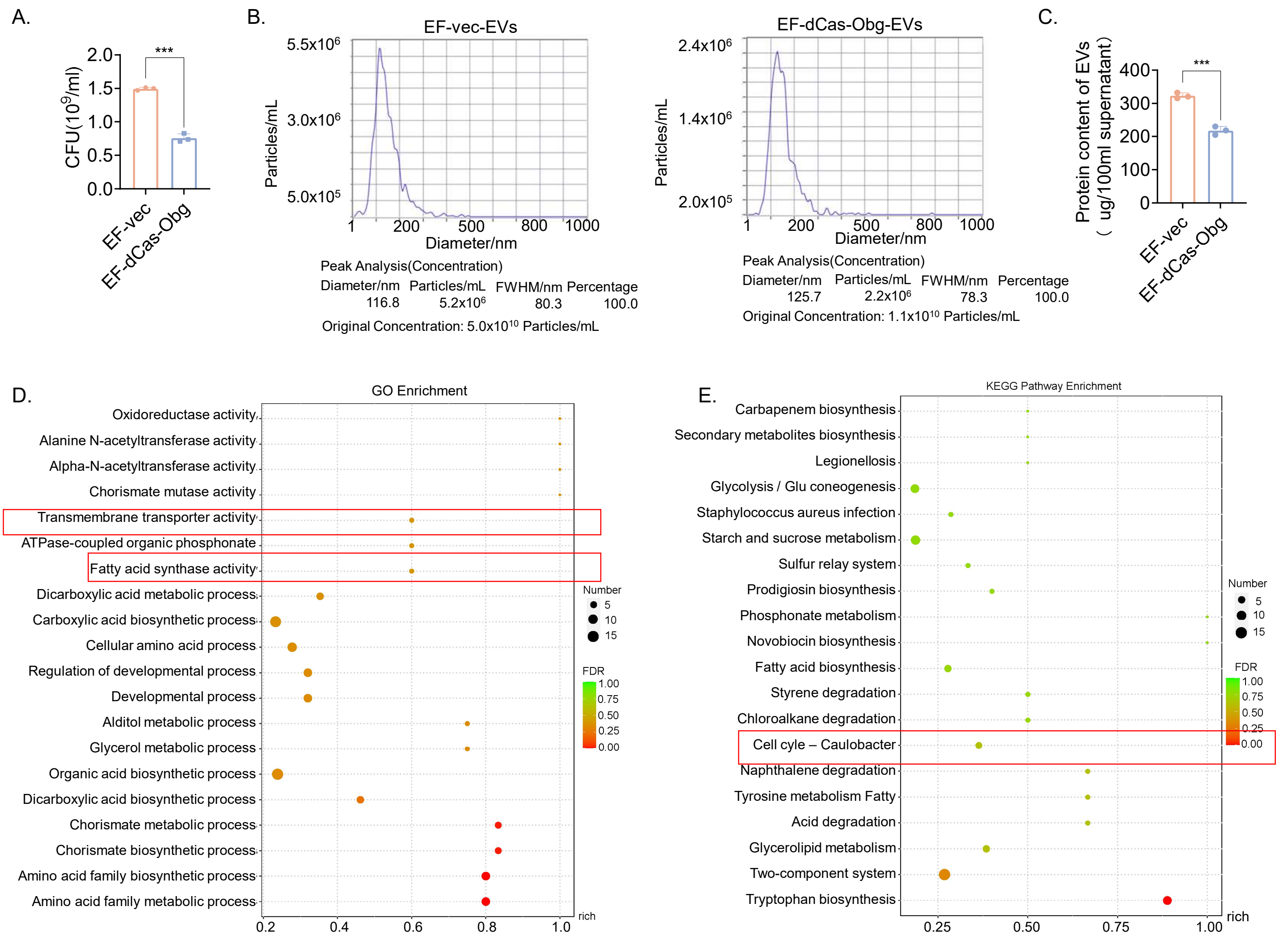

**Supplemental Figure 4 Engineered mutant strain of EF (Obg knock down strain, EF-dCas-Obg) compromised its growth rate, EVs production, and biological functions.**

(A) C.F.U calculation of EF-vec and EF-dCas-Obg strains after 12 hours cultured in BHI medium.

(B) Nanoparticle tracking analysis of EVs isolated from EF-vec and EF-dCas-Obg strains after normalizing with the total bacterial content.

(C) The protein concentration of EVs isolated from EF-vec and EF-dCas-Obg strains after normalizing with the total bacteria content.

(D) The gene ontology (GO) annotation results of the differentially expressed genes between EF-vec and EF-dCas-Obg strains.

(E) The Kyoto Encyclopedia of Genes and Genomes (KEGG) annotation results of the differentially expressed genes between EF-vec and EF-dCas-Obg strains.

Data are presented as mean ± SD; Unpaired two-tailed Student’s *t* test was used in (A), (C). ***, $p < 0.001$. C.F.U, Colony Forming Units.

Figure S5

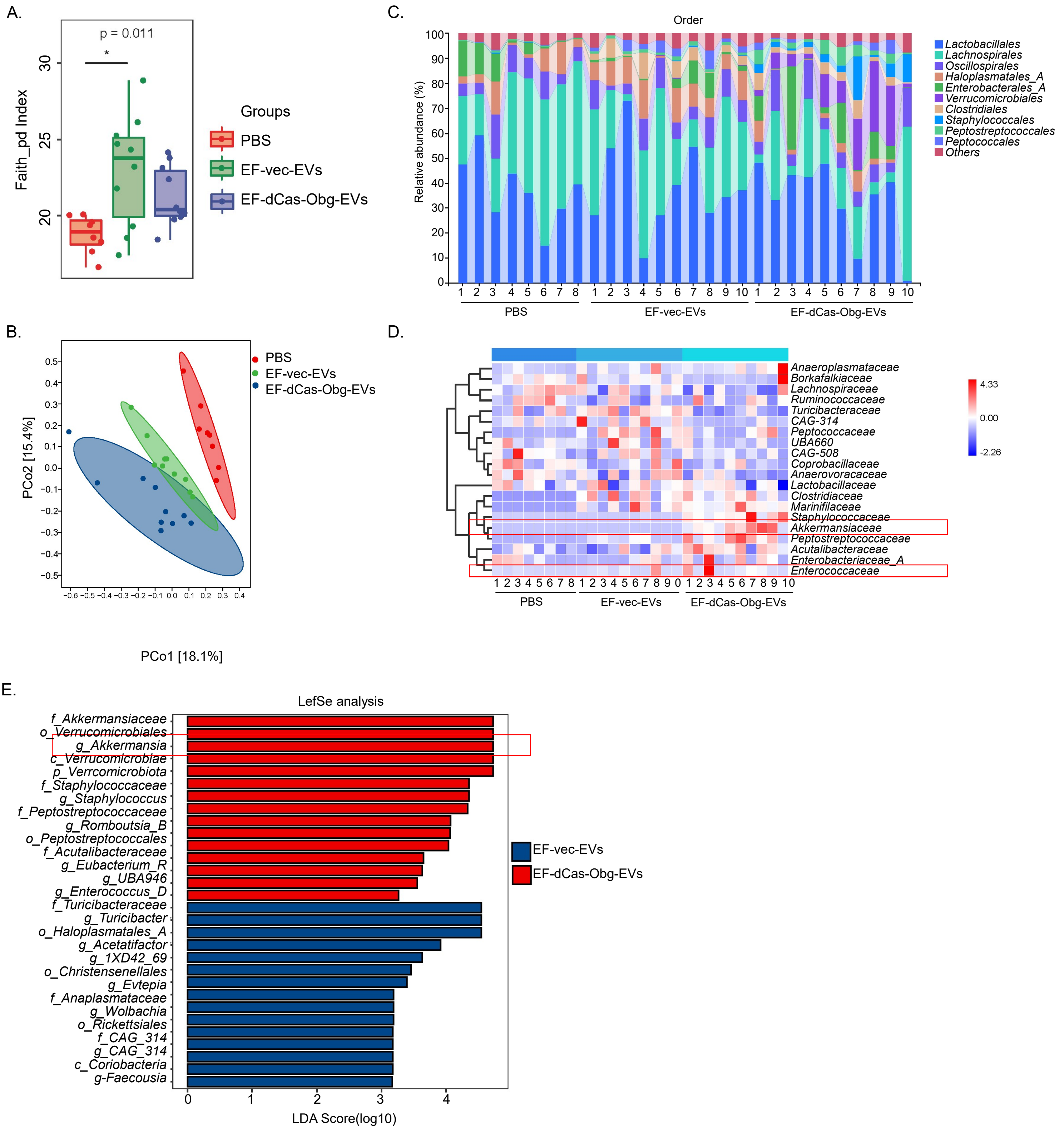

**Supplemental Figure 5. EF-dCas-Obg strain changed the composition of gut microbiome in mice**

(A) Faith_pd index showed the alpha diversity of microbiota in three groups.

(B) PCoA based on weighted UniFrac analysis of bacterial communities in three groups. Each dot represents a single sample.

(C) Relative abundance of mice fecal microbiota in three groups at Order level.

(D) Heatmap showing the distribution of various bacteria families in feces among 3 groups. The red boxes highlight altered bacteria *Akkermansiaceae* and *Enterococcaceae.*

(E) Linear discriminant analysis (LDA) effect size was used to identify the most differentially abundant taxa between two groups (LDA>3). The red boxes highlight increased *Akkermansia* genus.

Wilcoxon test was used to calculate the $p$ value in (A). One-way ANOVA test was used in (B). *, $p < 0.05$

Figure S6

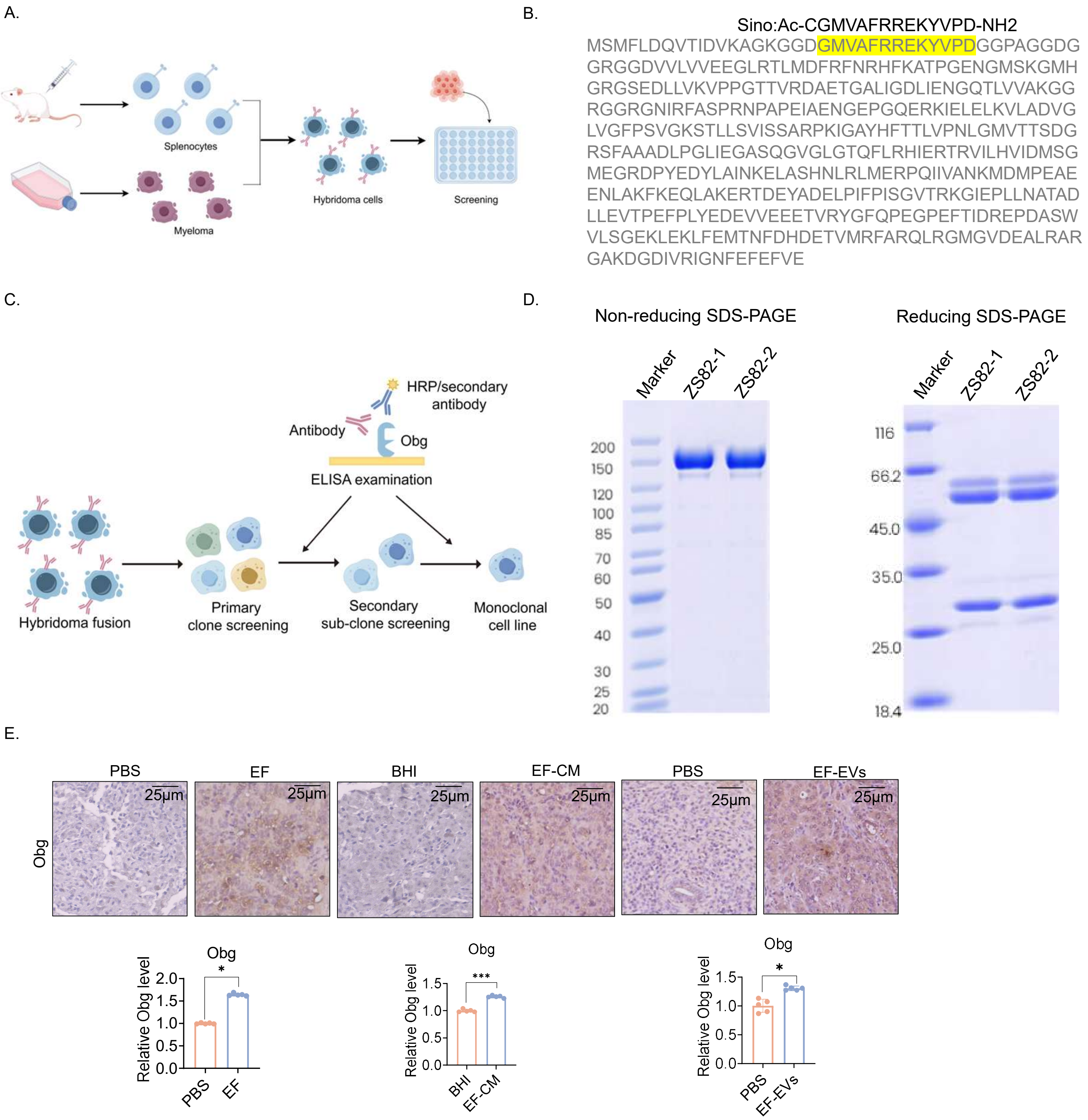

**Supplemental Figure 6 Preparation of anti-Obg antibody.**

(A) Schematic diagram of anti-Obg antibody production process.

(B) Peptide sequence aligned with Obg immunogen.

(C) Animal immunity and titer detection process.

(D) Non-reducing (left) and reducing (right) SDS-PAGE electrophoresis of purified antibody.

(E) Representative images of immunohistochemical staining of EF-Obg using the prepared anti-Obg antibody. Quantification of IHC staining was shown as bar graphs (below). Scale bar＝25μm.

Data are presented as mean ± SD; Unpaired two-tailed Student's *t* test was used in (E). *, $p < 0.05$; ***, $p < 0.001$.

**Supplemental Table 1: Association between EF-Obg expression and clinicopathological features in 100 HCC cases.**

| | Obg-low (n=30) | Obg-high (n=70) | *p* value |
|---|---|---|---|
| Gender | | | 0.425 |
| Female | 3 (10) | 4 (5.7) | |
| Male | 27 (90) | 66 (94.3) | |
| Age | 50.1±9.5 (31,72) | 52.1±13.0 (20,76) | 0.445 |
| AFP(ng/mL) | | | 0.965 |
| <25 | 14 (46.7) | 33 (47.1) | |
| ≥25 | 16 (53.3) | 37 (52.9) | |
| CEA(ng/mL) | | | 0.820 |
| <5 | 25 (83.3) | 57 (81.4) | |
| ≥5 | 5 (16.7) | 13 (18.6) | |
| CA199(U/mL) | | | 1.000 |
| <35 | 25 (83.3) | 59 (84.3) | |
| ≥35 | 5 (16.7) | 11 (15.7) | |
| Differentiation | | | 1.000 |
| Poor | 14 (46.7) | 34 (48.6) | |
| Moderate | 15 (50) | 34 (48.6) | |
| Well | 1 (3.3) | 2 (2.9) | |
| Tumor number | | | 0.044* |
| Single | 26 (86.7) | 47 (67.1) | |
| Multiple | 4 (13.3) | 23 (32.9) | |
| Tumor size(mm) | | | 0.048* |
| <50 | 21 (70) | 34 (48.6) | |
| ≥50 | 9 (30) | 36 (51.4) | |
| Macrovascular invasion | | | 0.497 |
| Absence | 28 (93.3) | 61 (87.1) | |
| Presence | 2 (6.7) | 9 (12.9) | |
| Microvascular invasion | | | 0.564 |
| Absence | 19 (63.3) | 40 (57.1) | |
| Presence | 11 (36.7) | 30 (42.9) | |
| Lymph | | | 0.088 |
| Absence | 28 (93.3) | 70 (100) | |
| Presence | 2 (6.7) | 0 (0) | |
| TNM stage | | | 0.035* |
| I | 15 (50) | 26 (37.1) | |
| II | 10 (33.3) | 24 (34.3) | |
| III | 3 (10) | 20 (28.6) | |
| IV | 2 (6.7) | 0 (0) | |
| BCLC stage | | | 0.013* |
| 0 | 6 (20) | 5 (7.1) | |
| A | 19 (63.3) | 37 (52.9) | |
| B | 1 (3.3) | 19 (27.1) | |
| C | 4 (13.3) | 9 (12.9) | |

The *p* values were calculated in SPSS19 using a chi-square test. All *p* values were two sided and the level of statistical significance was set at < 0.05. AFP, α-fetoprotein; CEA, Carcinoembryonic antigen; CA199, Carbohydrate Antigen 199.

### Supplemental Table 2: Key reagent and resource

| REAGENT or RESOURCE | IDENTIFIER | SOURCE |
|---|---|---|
| **Antibodies** | | |
| Anti-MYC Tag | GB12076 | Servicebio |
| Anti-GAPDH | 60004-1-Ig | Proteintech Group |
| Anti-HA Tag | 51064-2-AP | Proteintech |
| Anti-FLAG Tag | F1804 | Sigma |
| Anti-mTOR | 2983S | Cell Signaling Technology (CST) |
| Anti-Phospho-mTOR(Ser2448) | 5536S | Cell Signaling Technology (CST) |
| Anti-P70S6K | 14485-1-AP | Proteintech |
| Anti-Phospho-P70S6K(Thr389) | 9234s | Cell Signaling Technology (CST) |
| Anti-Vinculin | 4650s | Cell Signaling Technology (CST) |
| Anti-phospho-4EBP1(Ser65） | sc-293124 | Santa Cruz Biotechnology(SCBT) |
| Anti-HIF1α | GTX127309 | GeneTex |
| Anti-CD31(PECAM-1) | 77699 | Cell Signaling Technology (CST) |
| Anti-4EBP1 | 9644 | Cell Signaling Technology (CST) |
| Anti-Obg | N/A | This study |
| Anti-puromycin | ab315887 | Abcam |
| **Bacteria species** | | |
| *Enterococcus faecalis(ATCC 29212)* | N/A | Lab stock |
| *Enterococcus faecalis(OG1RF)* | N/A | Lab stock |
| *Escherichia coli BL(DE3)* | N/A | Lab stock |
| *Escherichia coli DH5α* | N/A | Lab stock |
| **Critical commercial assays** | | |
| E.Z.N.A. ® Universal Pathogen Kit | D4035-01 | OMEGA |
| Transcription/Translation Systems Kit | L1170 | Promega |
| Bacteria RNA Extraction Kit | R403-01 | Vazyme |
| FISH kit | D-0016 | EXONBIO |
| RT-PCR kit | 22948 | Promega |
| EasyPure HiPure Plasmid MiniPrep Kit | EM111-01 | TransGen Biotech |
| 10%(v/v) Cell Counting Kit-8 | K1080 | Apexbio Technology |
| **Chemicals** | | |
| Matrigel | 354248 | Corning |
| polybrene | TR-1003-G | Millipore |
| Cy7 | HY-D0825 | MCE |
| VivoGlo Luciferin | P1043 | promega |
| cytochalasin D | C102396 | Aladdin |
| dynasore | D423149 | Aladdin |
| Methyl-β-cyclodextrin | M413649 | Aladdin |
| Chlorpromazine | C424348 | Aladdin |
| isopropylβ-D-1-thiogalactopyranoside (IPTG) | I104812 | Aladdin |
| 3,3'-Dioctadecyloxacarbocyanine perchlorate | HY-D0969 | MCE |
| Everolimus | E125341 | Aladdin |
| Lipofectamine 2000 Transfection Reagent | 11668019 | Thermo Fisher |
| Nisin | HY-P1607 | MCE |
| Puromycin 2HCl | S7417 | selleck |
| Kanamycin | A100408-0005 | sangon biotench |
| Erythromycin | A600192-0025 | sangon biotench |
| Gentamicin | A100304-0001 | Santa Cruz Biotechnology |
| Ampicillin | A105483 | Aladdin |
| Vancomycin HCL | V105495 | Aladdin |
| 2×SYBR Green qPCR Master Mix | B21203 | BIMAKE |
| PrimeSTAR | R045B | Takara |

**Supplemental Table 3: Sequences for PCR**

| Primer | Sequence（5’-3’） |
|---|---|
| Obg-PCR | Forward: AGAATAGGAGGACAAATTATATGTCC<br>Reverse: GTGAACTGGTTTTATGCTTATTCGAC |
| (pCDNA3.1) HA-Obg-Myc | Forward: AAAGACGATGACGACAAGCTTATGTACCCTT<br>-ATGATGTGCCAGATTATGCCTCCATGTTTTTAGATCA<br>Reverse: CCACACTGGACTAGTGGATCCTTACAGATCTTC<br>-TTCAGAAATAAGTTTTTGTTCTTCGACAAATTCAAAT |
| (PET21a+) HA-Obg-Myc | Forward: CTCGAGTGCGGCCGCAAGCTTATGTACCCTTA<br>-TGATGTGCCAGATTATGCCTCCATGTTTTTAGATCA<br>Reverse: CAGCAAATGGGTCGCGGATCCTTACAGATCTT-<br>CTTCAGAAATAAGTTTTTGTTCTTCGACAAATTCAAAT |
| (pLVX-Puro) HA-Obg-Myc | Forward: TCGAGCTCAAGCTTCGAATTCATGTACCCTTAT-<br>GATGTGCCAGATTATGCCTCCATGTTTTTAGATCA<br>Reverse: GTACCGTCGACTGCAGAATTCTTATTCGACAA-<br>ATTCAAATTCAAAATT |

**Supplemental Table 4: Target EF-Obg sequences predicated by CHOPCHOP**

| Target sequence | Efficiency |
|---|---|
| GGTATGAGTAAAGGAATGCACGG | 68.36 |
| TCACGAACTGTTGTACCTGGTGG | 67.60 |
| GGCACCAATTTTCGGACGAGCGG | 64.15 |

**Supplemental Table 5: Primer sequences for RT-qPCR**

| Primer | Sequence（5’-3’） |
|---|---|
| Total bacteria-16S | Forward: GCAGGCCTAACACATGCAAGTC<br>Reverse: CTGCTGCCTCCCGTAGGAGT |
| Obg | Forward: GAACCGGGTCAAGAACGTAAA<br>Reverse: GTCCTTCCATGCCACTCATATC |
| *E.faecalis* | Forward: CGCTTCTTTCCTCCCGAGT<br>Reverse: GCCATGCGGCATAAACTG |
| *E.casseliflavus* | Forward: GGAAGAAAGTTGAAAGGC<br>Reverse: TCGGTCAGACTTKCGTCC |
| *E.faecium* | Forward: TGCTCCACCGGAAAAAGA<br>Reverse: CACCAACTAGCTAATGCA |
| *E. mundtii* | Forward: ATTCGATTCCCTGAGTAGCG<br>Reverse: TCCAAATTTCATCTACGGGG |
